# Why Do Hybrids Turn Down Sex?

**DOI:** 10.1101/2023.02.24.529842

**Authors:** Frederic Fyon, Waldir Miron Berbel-Filho, Ingo Schlupp, Geoff Wild, Francisco Ubeda

**Affiliations:** Royal Holloway University of London; University of Oklahoma; University of Western Ontario; Royal Holloway

**Keywords:** Asexual Reproduction, Maintenance of Sex, Evolution of Asexuality, Hybridization, Reproductive Assurance, Amazon Molly

## Abstract

Asexual reproduction is ancestral in prokaryotes; the switch to sexuality in eukaryotes is one of the major transitions in the history of life. The study of the maintenance of sex in eukaryotes has raised considerable interest for decades and is still one of evolutionary biology’s most prominent question. The observation that many asexual species are of hybrid origin have led some to propose that asexuality in hybrids results from sexual processes being disturbed because of incompatibilities between the two parental species’ genomes. This proximate theory appears difficult in real life, as it requires fundamental reproductive traits to be profoundly altered without collapsing individuals’ fertility. Repeated failures to produce asexual Amazon Molly in the lab through crossing experiments show that we are still in need of an evolutionary explanation. Here, we present a mathematical model and propose an adaptive route for the evolution of asexuality from previously sexual hybrids. Under smaller reproductive alterations, we show that asexuality can evolve to rescue hybrids’ reproduction. Importantly, we highlight that when incompatibilities only affect the fusion of sperm and egg’s genomes, unreduced meiosis and paternal genome elimination can evolve separately, greatly facilitating the overall evolutionary route.

## INTRODUCTION

Sexual reproduction whereby individuals produce gametes – carrying one copy of each gene – that combine with other gametes to produce a new individual – carrying two copies of each gene, one from each parent –, is the dominant form of reproduction in organisms formed by cells with a nucleus (Eukaryotes). Sexual reproduction evolved from asexual reproduction in what has been considered a major transition in evolution [1]. In spite of abundant research in the topic, the drivers of this major transition remain debated. Although sexual reproduction in Eukaryotes is stable over long periods of evolutionary time, certain forms of asexual reproduction have in turn evolved from sexual reproduction multiple times [2, 3, 4]. Some of the taxonomic groups where asexual reproduction has evolved independently include anemons, nematods, molluscs, arthropods, and importantly vertebrates (see Fig 1 and references therein).

**Figure 1:**
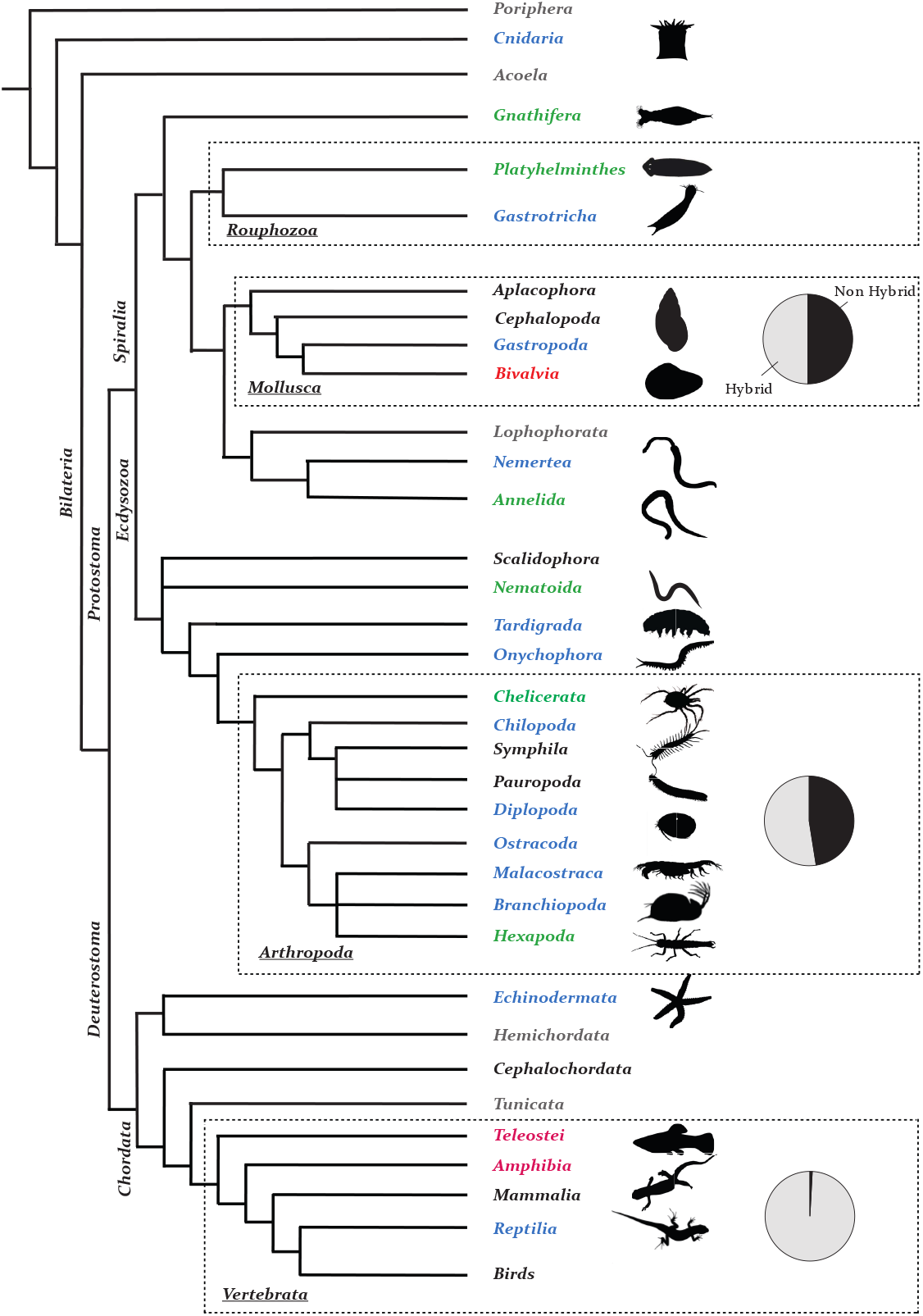
Distribution of reproductive modes within a simplified phylogeny of metazoans (adapted from [5, 6, 7]). We are aware that some of the phylogenetic relationships represented here are debated, and that we are comparing taxa at different levels; this phylogeny is presented merely for the purpose of illustrating the broad distribution of the different sexual and asexual reproductive modes among metazoans. Taxonomic groups for which there has been reports of true parthenogenetic species or races appear in blue, of gynogenetic (sperm-dependent parthenogenetic) species or races appear in red, and of both types of species or races appear in green. Grey taxa correspond to groups where vegetative reproduction (budding, fission) has been reported, while all species in black taxa exclusively reproduce sexually. Dashed rectangles highlight clusters of taxa where parthenogenesis (true or sperm-dependent) occurs more often: flat and spined worms (*Rouphozoa*), molluscs, arthropods and vertebrates. On the right, we added profiles of some asexual species corresponding to the colorized taxa. Further on the right, we present as pie charts the proportion of asexual species that are of hybrid origins (grey) compared with other origins of asexuality (black) based on [3]. While molluscs and arthropods show approximately half of asexual species being of hybrid origin, vertebrate asexuals are most if not all of hybrid origin.

It is not clear how asexuality has evolved so many times in such diverse taxa. On one hand, the evolutionary advantages that drove sexuality to outcompete asexuality originally are expected to still apply in these taxa. On the other hand, empirical evidence shows that asexual species evolving from sexual ancestors have to overcome many cytological and physiological obstacles [8, 9]. In spite of the abundant scientific interest on what factors may drive the transition from sexuality to asexuality, these drivers remain elusive [3].

One intriguing observation is that many of the asexual species evolving from sexual ancestors originated from the cross of two different species, i.e. are of hybrid origin [3, 10, 11] (Fig 1). Of those surveyed, half of all asexual species in molluscs and arthropods, and all but one asexual species in vertebrates are of hybrid origin [3, 12] (Fig 1). Why does hybridization favour the evolution of asexuality? Understanding the drivers of this transition will help us understand the evolutionary stability of sexual reproduction, a fundamental feature of multicellular organisms’ physiology, behaviour, diversity and evolution.

It has been argued that asexuality can appear as the direct outcome of crossing two different sexual species. Incompatibilities between genes of different species (*genomic incompatibilities*) are thought to be able to disrupt key processes in sexual reproduction, such as meiosis and/or gamete recognition, leading to the spontaneous production of fully functional asexual progeny [13, 14, 15]. Crossing experiments provide partial support as they show that it is possible to obtain asexual progeny spontaneously in some species [15, 16, 17, 18] but not in others [19, 20, 21, 22, 23]. Here we explore the conditions for natural selection to favour the evolution of asexuality from sexuality in those cases where asexuality does not appear spontaneously as a direct result of genomic incompatibilities. In doing so, we will explore whether a progressive evolution of asexuality is possible and whether hybridization may act as a catalyst of the transition to asexuality.

We chose as model for our study a well-established example of crossing experiments that do not result in the spontaneous production of asexual progeny: the Amazon Molly (*Poecilia formosa*). The Amazon Molly is a species of female-only live-bearing fish from coastal streams of the Gulf of Mexico. It originated at least 100,000 years ago from the hybridization between Sailfin Molly (*Poecilia latipinna*) males and Atlantic Molly (*Poecilia mexicana*) females [23, 24]. Amazon Molly females reproduce asexually by producing diploid clonal eggs whose development is activated by fusion with sperm of parental species even though the paternal genetic material is lost [10, 11]. Thus the progeny of Amazon Molly is diploid and clonal [25]. This form of asexuality is known as gynogenesis and, although relatively rare, it has been reported in most taxa where other forms of asexuality has been observed [17] (Fig 1). Laboratory crosses between modern Sailfin and Atlantic Mollies have failed to produce asexual gynogenetic hybrids multiple times [19, 20, 21, 22, 23]. Instead, females and male hybrids progeny from experimental crosses are sexual. Further support for the sexuality of the hybrid progeny of Sailfin and Atlantic Mollies comes from the genome of the Amazon Molly itself. This genome carries the signature of crosses between Amazon Molly’s hybrid ancestors and their parental species [26]. The adaptive forces that favoured the evolution of modern asexual Amazon Mollies from hybrid sexual ancestors remain unknown.

For the spontaneous appearance of gynogenetic hybrids from the cross of Sailfin and Atlantic Mollies – and in general for the appearance of any gynogenetic hybrid, two reproductive processes would need to be disrupted. First, the reduction in ploidy of germ cells – from diploid to haploid – would need to be prevented to generate diploid gametes (*unreduced meiosis*). Second, the fusion of maternal and paternal pronuclei in eggs fertilised by sperm would need to be averted to impede the transmission of the paternal genome (*paternal genome elimination*). In addition, all progeny should consist of females as gynogenetic species are female-only. It has been argued that it should be difficult, and thus rare, for genetic incompatibilities to result in the modification of all these aspects of the reproductive systems at once to produce a viable and fertile asexual progeny [11]. Instead, it has been suggested that a progressive modification of the reproduction of the hybrids would be much easier evolutionarily, and should thus account for more transitions from sexuality to gynogenesis [11]. However, this verbal argument has not been backed by any formal model. Here we formulate a novel mathematical model for the evolution of asexual gynogenesis from sexual reproduction one step at a time. We model the ecological conditions and the sequence of intermediate steps that may lead to modern Amazon Mollies.

While using the well-researched case of the Amazon Molly as a reference, to help us contextualise our results, our evolutionary mathematical model applies to any other asexual gynogenetic species. Furthermore our conclusion can be extended to other forms of asexuality beyond gynogenesis. In this research we show that natural selection favours the progressive (one step at a time) evolution of asexuality: for the first time, we propose an evolutionary route that decouple the evolution of unreduced meiosis and paternal genome elimination. The reason why each of the incremental steps are favoured by natural selection is that each of them increases the reproductive opportunities of hybrids with otherwise limited reproductive opportunities (i.e. increase *reproductive assurance*). We thus provide progressive routes that are plausible alternatives to the restrictive spontaneous route that currently prevails as an explanation. Furthermore, we argue that these progressive routes are only available to hybrids, thus explaining why hybrid species are hotspots for the transition from sexuality to asexuality.

## METHODS

Here we model the interactions between two species (*P. mexicana* and *P. latipinna*) and the hybrid resulting from crosses between them. We refer to *P. mexicana* and *P. latipinna* as the *parental species* and the hybrid they produce as the *wild-type hybrid*.

Parental species and hybrids are assumed to coexist within a same environment. There has been different theories proposed to explain how a gynogenetic hybrid can coexist with its parental species while being a resource and mate competitor ([27, 28, 29, 30]). In this work, we use the simplest of this assumption, and extend it to sexual and asexual hybrids: we assume that parental species and hybrid population do not compete for resources. That is, the three populations coexist in a given environment with death rates Ψ_*i*_ that depend only on the total number of individuals of that species *i*: Ψ_*i*_ = 2*N_i_/K* (*N_i_* representing half the number of individuals from species *i* present in the environment, see Supplementary Material for further justification). Henceforth, we use subscripts *m* for the *P. mexicana* species, *l* for the *P. latipinna* species, and *h* for the hybrid species. Because we assume perfect symmetry between the parental species, we will generally use subscript *p* to refer to any of the two parental species. To simplify equations, we will use in the following *n_i_*, the population sizes relative to the carrying capacity of the environment (*n_i_* = *N_i_/K*).

Birth rates are influenced by the mating preferences of the different populations (illustrated on Fig 2). We note *c_p_* the preference of a parental female (*mexicana* or *latipinna*) for males from its own population, and *c_h_* the preference of hybrid females for hybrid males.

**Figure 2:**
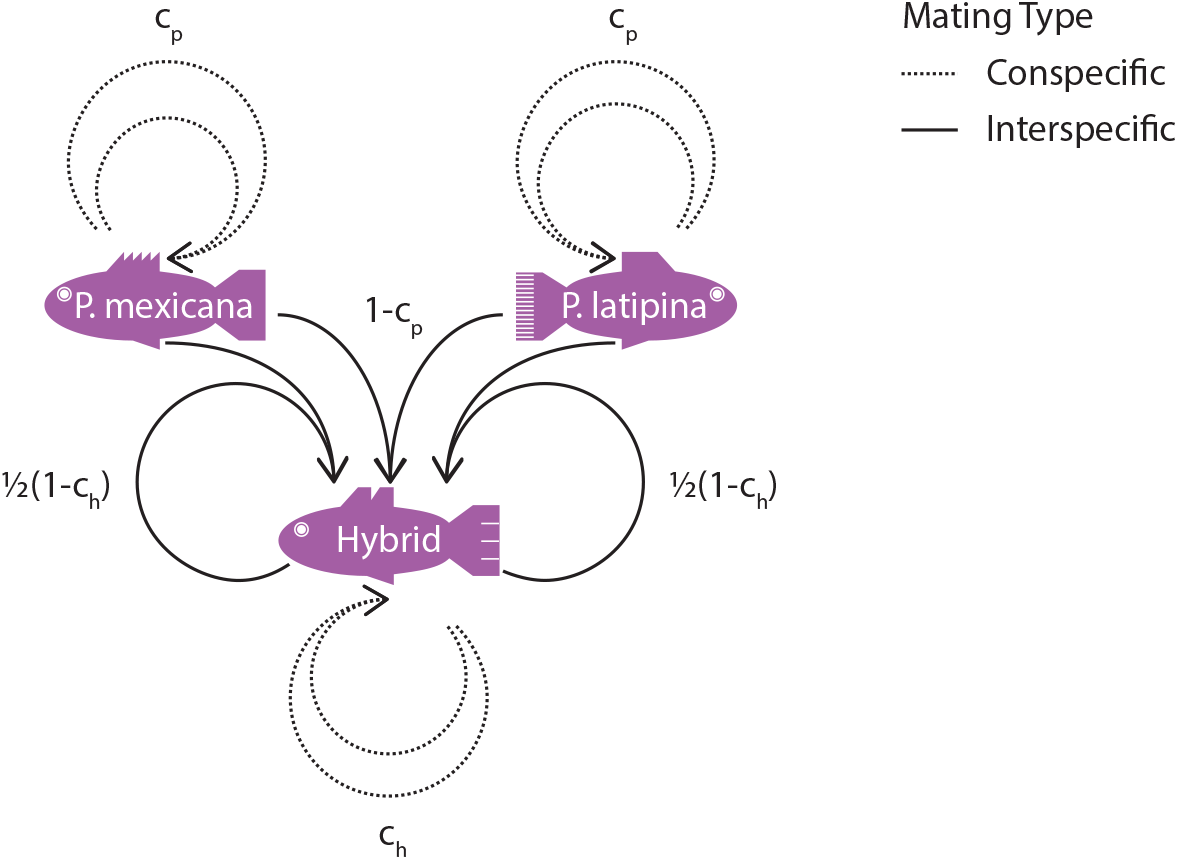
Reproduction and mating choices. Females from each parental species (*P mexicana* on the left and *P. latipinna* on the right) mate with males from the same species with preference *c_p_* (Conspecific Matings), which gives birth to individuals of the same species. Alternatively, they can choose to mate with a male from the other parental species with preference 1 – *c_p_* (Interspecific Matings); in this case, they give birth to hybrid individuals. Females from the hybrid species can choose to mate with hybrid males with preference *c_h_* (Conspecific Matings), or with males from the parental species with preference 1 – *c_h_* (Interspecific Matings). In both cases they give birth to hybrid individuals. Conspecific Matings are showed in dashed lines, while Interspecific Matings are shown in plain lines.

As a result of birth by intra-specific matings and death by intra-specific competition for resources, the parental populations reach a stable equilibrium population size 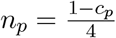 (see Supplementary Material for a demonstration). Henceforth, we will always assume that the parental population size is equal to this stable equilibrium.

Wild-type hybrid females mate with hybrid males with probability Φ. Φ is an increasing function of *n_h_* and *c_h_* whose expression is given in the Supplementary Material: Φ increases when hybrids female preference for hybrid males increases, or if there are relatively more hybrid males around. Note that hybrid and parental species do compete for mates; thus Φ also depends on the amounts of paternal males around, that is, on *c_p_* (Φ decreases if there are relatively more parental males around). Alternatively, wild-type hybrids can mate with parental males (back-crossing) with probability 1 – Φ. Both kinds of matings, when productive, produce wild-type hybrids (see Fig 2).

The size of the population of wild-type hybrid females changes as a result of the birth and death mechanisms described above following:

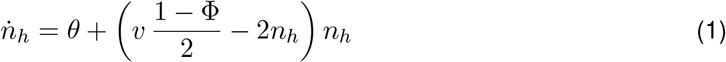

with *v* denoting the viability of offspring produced by sexual hybrid females when backcrossing with parental males. We will see later on that we assume that this value is either 0 or 1, depending on the model of genomic incompatibilities we consider. *θ* refers to the birth of wild-type hybrids by hybridization between the parental species: *θ* = *n_p_*(1 – *c_p_*) = *c_p_*(1 – *c_p_*)/4.

It is easy to show that the dynamics in Eq 1 lead the wild-type hybrid female population to a stable equilibrium size 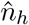 (see Supplementary Material for a demonstration). Depending on the model of genomic incompatibilities we consider (see below), we can in some cases find an analytical closed-form expression of 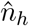. In all cases, we can find 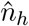 at least numerically.

We generally assume that both parental species and the wild-type hybrid population are fully sexual. Once the wild-type hybrid has reached its equilibrium, we introduce in the environment a *mutant hybrid* that can potentially display any mode of reproduction between fully sexual and fully asexually gynogenetic. In particular, the phenotype of the mutant hybrid species is characterised by three evolutionarily labile traits: production of diploid clonal eggs with probability *α* ∈ [0, 1]; elimination of the paternal genome with probability *β* ∈ [0, 1], and production of female progeny in proportion *σ* ∈ [-1, 1] (the proportion of females relative to males in the progeny (sex-ratio) is 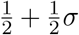). Importantly, these are female-only phenotypes; mutant hybrid males do not express *α*, *β*, or *σ* though they do carry the genes for each. Notice that the phenotype corresponding to the hybrid wild-type species is (*α* = *β* = *σ* = 0) and that of the modern Amazon Molly is (*α* = *β* = *σ* = 1).

Of the three traits above, let’s focus for a moment on the mutant trait *α* and consider the different ways an organism can produce diploid clonal eggs. There are two ways in which diploid clonal eggs can be produced [8, 31]. First, they can be produced by circumventing meiosis to produce gametes via mitotic (unreduced) division, a process referred to as *apomixis*. Alternatively, they can be produced by duplicating the genome prior to meiosis to produce gametes via meiotic (reduced) division, a process referred to as *endoduplication*. In the latter case, meiotic products are identical to the germ-cell they originated from because recombination occurs between identical sister chromatids and therefore does not produce new combinations of genes. Because Amazon Mollies produce clonal eggs through apomixis [32], henceforth we will just use the term apomixis to refer to clonal egg production but our model applies to endoduplication – considered to be the mechanism used by many other gynogenetic vertebrates [33].

Our model assumes that incompatibilities between parental species lead to some form of egg or sperm dysfunction in hybrids, which in turn affects the reproductive prospects of female hybrids.

We first consider the case where hybrid sperm is viable but unable to decondense its own pronucleus (henceforth the *sperm-fails-to-decondense scenario*). Thus, while all sperm is able to trigger embryogenesis, hybrid sperm is unable to contribute genetic material to the embryo while parental sperm is (see Figure 3, top panel). In this case, meiotic hybrid females can produce viable diploid offspring by mating with parental males and incorporating the paternal genome (*v* = 1). Apomictic hybrid females, however, can produce viable diploid offspring by either discarding the paternal genome or by mating with hybrid males (see Figure 3, top panel, right-hand side). Notice that throughout this research we assume that haploid and triploid offsprings are lethal for simplicity. Empirical work shows that Amazon Molly triploids have reduced fitness even though they fall short of being lethal [34]. Notice also that, when mating with hybrid males, mutant hybrid females can achieve greater success through apomixis than via meiosis.

**Figure 3:**
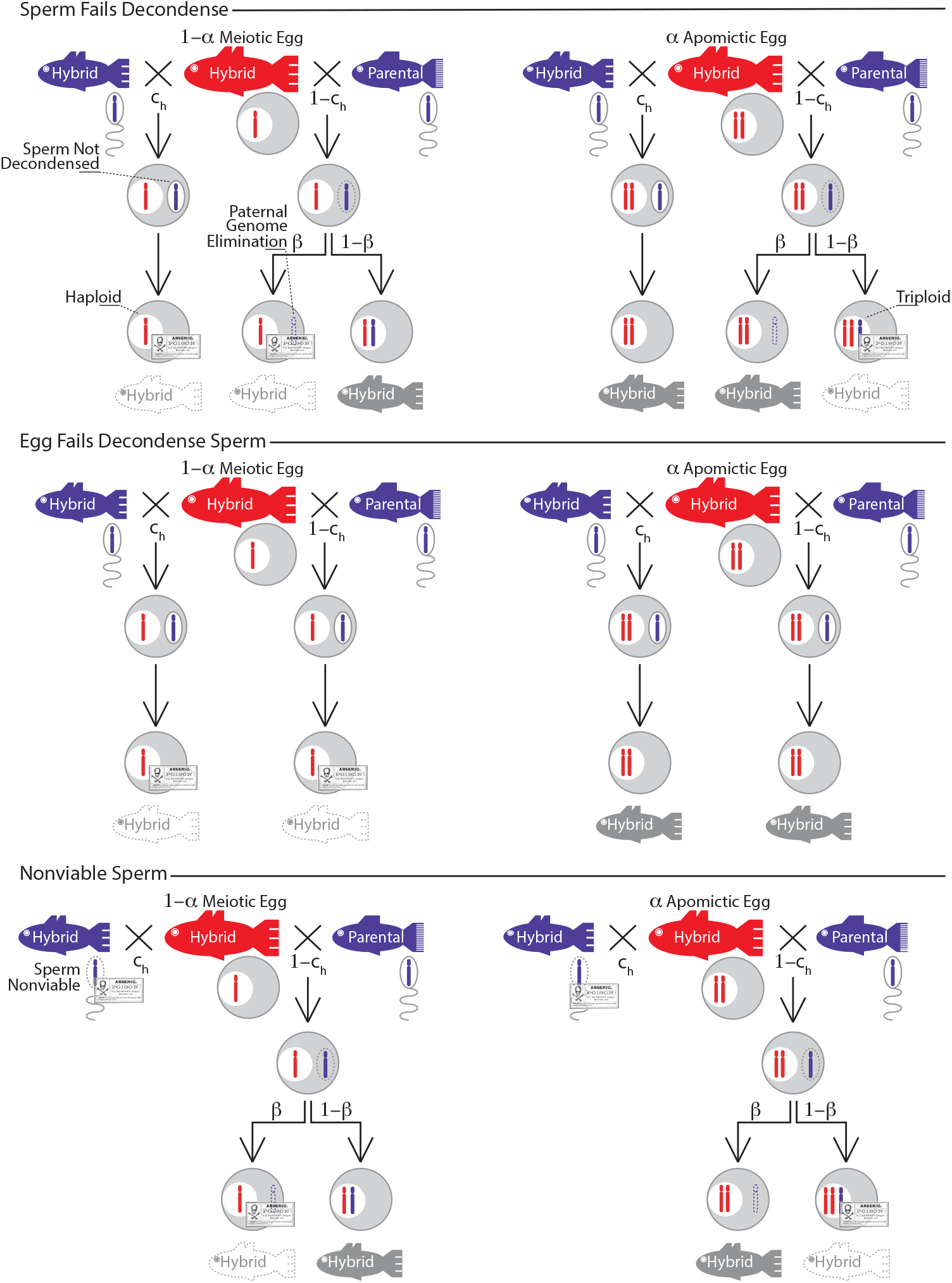
Hybrid female reproduction depending on the model considered, on the type of mate (hybrid or parental), on the type of egg (meiotic or apomictic, the latter occurring with probability *α*) considered, and on the occurrence of paternal genome elimination (with probability *β*). Unviable offsprings are indicated with crossbones and white, dashed-bordered fishes. This happens whenever the embryo is haploid or triploid. The mother genetic naterial is indicated in red, while the father genetic material is shown in blue. Grey-filled fishes indicate viable offspring. where *v* (already mentioned), *s* and *x* allow to take into account the differences induced by the assumptions of the three scenarios on the growth rate of mutant hybrids. In the *sperm-fails-to- decondense scenario*, we have *v* = 1, *s* = 1 and *x* = 1. In the *egg-fails-to-decondense-sperm-scenario*, we have *v* = 0, *s* = 1 and *x* = 0. Finally, in the *nonviable-sperm-scenario*, we have *v* = 1, *s* = 0 and *x* = 1. See Supplementary Material for the corresponding equations with *v*, *s* and *x* replaced by their values in each scenario.

Second, we consider the alternative case when hybrid eggs are unable to decondense the pronucleus of the sperm fusing with the egg (henceforth *egg-fails-to-decondense-sperm scenario*). Thus, all sperm is able to trigger embryogenesis but unable to contribute genetic material to the embryo (see Figure 3, middle panel). In this case, meiotic hybrid females cannot produce viable diploid offspring with any type of males (including parental males: *v* = 0). Apomictic hybrid females however, can produce viable diploid offsprings with any type of males (see Figure 3, middle panel, right-hand side). This time, mutant hybrid females can achieve greater success through apomixis than they can with meiosis when mating with any kind of males.

Finally, we consider the case when hybrid sperm is nonviable (henceforth *nonviable-sperm scenario*). Now, hybrid sperm is unable to trigger embryogenesis and unable to contribute genetic material to the embryo while parental sperm is able to perform both functions (see Figure 3, bottom panel). In this case, hybrid females (both meiotic and apomictic) can only produce viable offspring if they mate with parental males (*v* = 1). That said, mutant hybrid females employing meiosis must incorporate the paternal genome to produce viable offspring, whereas those employing apomixis must eliminate the paternal genome to produce viable offspring.

Once a rare mutant hybrid female has been introduced in the environment, it should spread and invade if and only if it is able to produce more than one mutant hybrid daughter during its lifetime. Mathematically, this means that their growth rate, that we can also call fitness *w*, is strictly greater than 0. We can find a general equation for *w* that works for the three models:

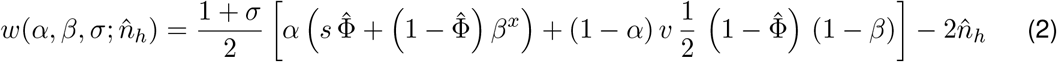

If a mutant hybrid invades, then it coexists with the parental species and wild-type hybrids, until such time as a new mutant hybrid arises. If this new mutant has a selective advantage (resp. disadvantage) relative to the established one, the new mutant will eliminate (resp. be eliminated by) its predecessor and establish itself (resp. be absent) in the equilibrium ecosystem. Ultimately, hybrid traits evolve in the long term through a succession of mutant invasion and displacement events. Because the equilibrium size of the mixed population of wild-type and established mutant hybrids is independent of the traits expressed by any new mutant that may arise, in the long-term selection will act to maximize *w*. Thus, by considering where *w* is positive and how it can be maximized, we seek to capture the transition from an ancestral sexual hybrid to a hybrid whose phenotype more closely matches that of the modern Amazon Molly.

Finally, we carry out simulations to establish the order in which this transition may have occurred over evolutionary time. We assume that evolution begins with the invasion of a mutant with the lowest values of *α*, *β* and *σ* such that 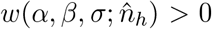. We numerically calculate the resulting wild-type - mutant equilibrium. Then, we calculate selection gradients by taking the partial derivatives of 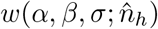 with respect to *α*’, *β*’ and *σ*’. Here, *α*, *β* and *σ* refer to the established mutant’s phenotype, while *α*’, *β*’ and *σ*’ refer to a new, rare mutant. Note that here 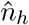 refers to the total number of hybrid females at the wild-type - mutant equilibrium. By comparing these and finding the highest derivative, we find the direction towards which selection is pointing at the current wild-type - mutant equilibrium. We assume that a mutant with a phenotype marginally different in that direction then spreads, invades, replaces the older mutant and we calculate the new wild-type - mutant equilibrium. We repeat this process until no new mutant can spread. The algorithm used for these numerical simulations is available in the Mathematica file provided as supplementary material.

## RESULTS

### Sperm-Fails-to-Decondense Scenario

We start by considering whether a rare mutant that deviates from the wild-type reproductive phenotype, i.e. *α* = *β* = *σ* = 0, can invade the hybrid population. That is, we set to find out the conditions for 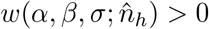. In the case of this first model (*v* = 1, *s* = 1 and *x* = 1), this means:

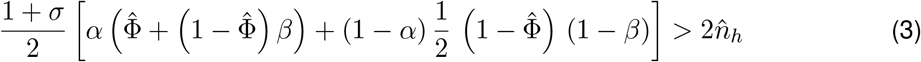

Using the fact that 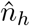 and 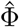 are by definition such that *ṅ_h_* = 0, Eq 1 allows us to show that a necessary condition for 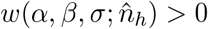 is *α* > 0 (see Supplementary Material for a more detailed demonstration). Importantly, however, neither *β* = 0 nor *σ* = 0 preclude mutant invasion. Thus, in this scenario, a mutant must necessarily display some strictly positive rate of production of apomictic eggs to have a chance to invade.

If a mutation can only modify one phenotypic trait, we understand that the first mutant to invade must be partially apomictic. To obtain the invasion condition of such a mutant, we can thus assume *β* = *σ* = 0 to simplify the equations. Replacing 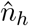 by its expression in function of *θ* and 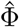 obtained by solving *ṅ_h_* = 0, this leads to:

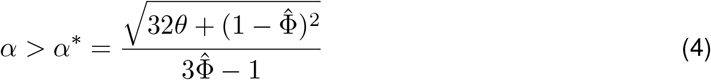

with *α*\* being the lower bound of *α* such that a mutant with phenotype (*α*, 0, 0) can invade.

We plot on Fig 4.A *α*\* in function of *c_h_* and *c_p_*, calculating 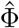 numerically. *α*\* decreases with the preference for hybrids to mate with other hybrids *c_h_*. In the absence of paternal genome elimination, apomictic hybrid females gain reproductive opportunities with hybrid males, but lose reproductive opportunities with parental males (see Fig 3, top panel, right-hand side). Thus, mutant partially apomictic females gain a reproductive advantage over wild-type meiotic females only when the probability of hybrid females mating with hybrid males 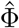 is large enough; probability that increases with the preference of hybrid females for hybrid males *c_h_*.

**Figure 4:**
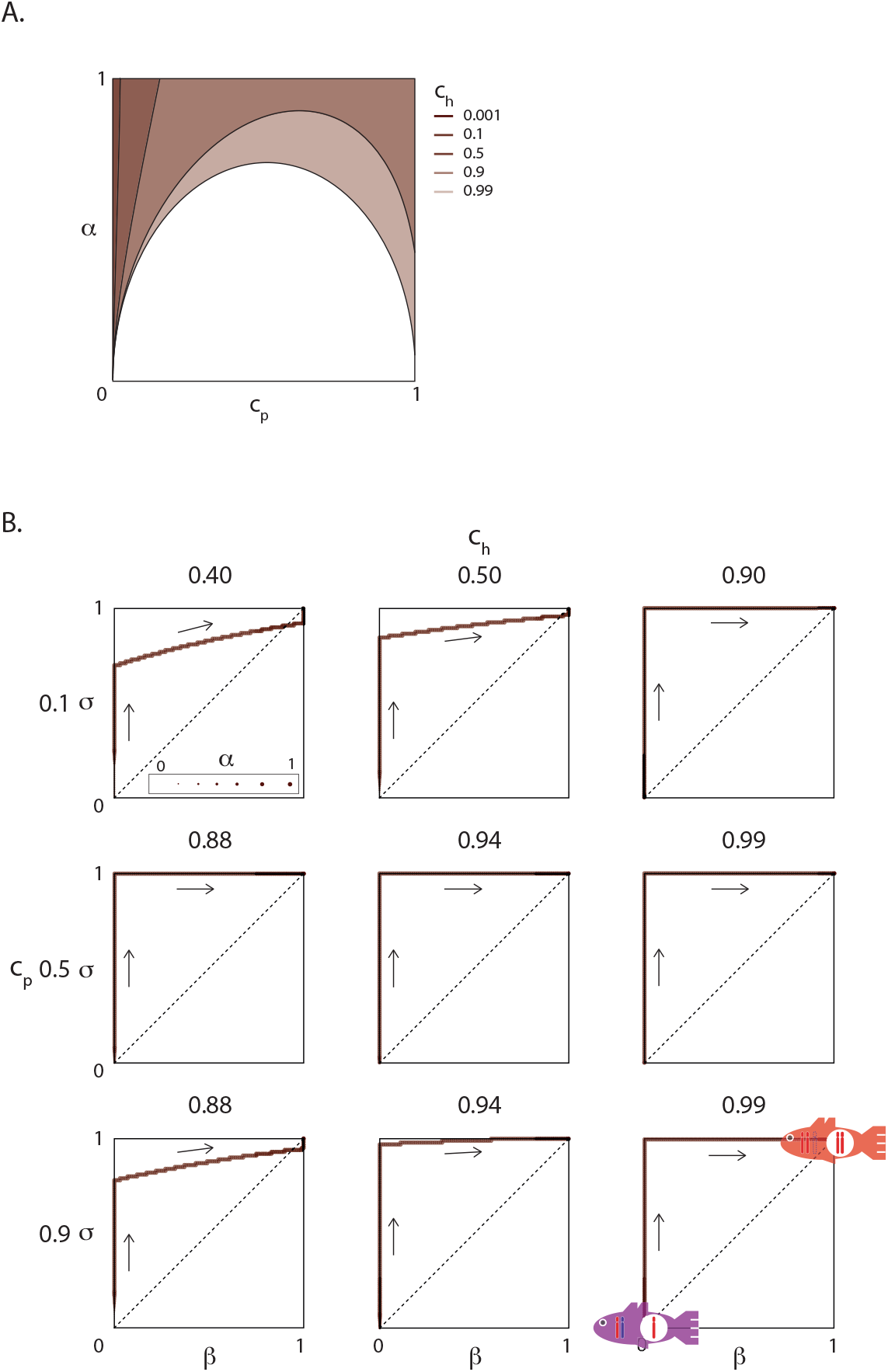
Evolution of gynogenesis in the *sperm-fails-to-decondense scenario*. A. Values of *α* > *α*\* (y-axis) where a mutant can invade are indicated in function of *c_p_* (x-axis) and *c_h_* as shaded areas. Lighter colours indicate growing values of *c_h_*. B. Predicted evolutionary route to gynogenesis for nine combinations of *c_p_* and *c_h_* values. The size of the dot represents *α*, while x- and y-axes represent *β* and *σ* respectively. These routes show the transition from sexual hybrids (purple fish), which are diploid, produce haploid eggs and have maternally (red) and paternally (blue) inherited chromosomes, to gynogenetic hybrids (red fish), which are diploid, produce diploid eggs and have only maternally-inherited chromosomes. The bottom limit *α*\* of *α* for the first mutant to spread varies in each case, and approximately equals respectively from left to right and top to bottom: (*c_p_* = 0.1) 0.89, 0.72, 0.62; (*c_p_* = 0.5) 0.91, 0.80, 0.72; (*c_p_* = 0.9) 0.45, 0.59, 0.80.

*α*^*^ reaches a maximum at intermediate values of the preference of each parental species to mate with the other parental species *c_p_*. This is because *θ*, the influx of wild-type hybrids by direct hybridization, is proportional to *c_p_*(1 – *c_p_*). *θ* is maximal at intermediate values of *c_p_*, which increases *α*\* as can be seen from Eq 4. Larger values of *θ* means larger values of *n_h_*. The invasion of a mutant is thus compromised by large values of *θ* (and thus intermediate values of *c_p_*) as it tends to increase the death rate of mutant hybrids due to competition for resources with increased numbers of wild-type hybrids.

As explained in the Methods section, we can find the long-term evolutionary route of this system by derivating 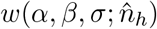, that is by looking at fitness gradients. We plot on Fig 4.B these evolutionary routes. It stands out from these numerical computations that if the conditions are such that a first, partially apomictic mutant can invade (*α* > *α*\*), then gynogenesis (*α* = *β* = *σ* = 1) ends up evolving always. We show in the Supplementary Material that indeed gynogenesis ultimately maximizes the mutant fitness function. Also, we see on Fig 4.B that most of the time the evolution of gynogenesis happens in a stepwise process, with first the evolution of females producing exclusively haploid eggs (*α* = 1), then the evolution of fully female-biased sex-ratios within the offspring (*σ* = 1), and finally the evolution of females always eliminating paternal genomes (*β* = 1). Most importantly, these evolutionary routes show that the evolution of apomixis occurs before and independently from the evolution of paternal genome elimination. In restricted parameter spaces, there may be some limited overlap between these steps, but this does not affect the qualitative prediction that in this model the evolution of apomixis does not require the evolution of paternal genome elimination in parallel.

In the following, we follow the same analysis for the other two models.

### Eggs-Fails-to-Decondense-Sperm Scenario

From a modelling perspective a failure of hybrid eggs to decondense sperm is equivalent to assuming that paternal genome elimination is a direct result of hybridization. Therefore in this scenario, the wild-type hybrid has a reproductive phenotype characterised by (*α* = 0, *β* = 1, *σ* = 0). Again, we start by considering whether a rare mutant that deviates from the wild-type reproductive phenotype can invade the hybrid population.

In this scenario (*v* = 0, *s* = 1 and *x* = 0), the mutant invasion condition 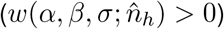 is:

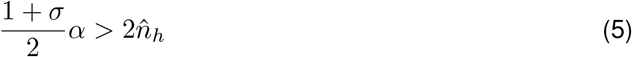

Because 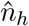 is necessarily positive, we realize that again, invasion can only happen for values of *α* strictly positive. Because this is not the case for *σ*, in this scenario the first mutant to possibly invade must again be a partially apomictic mutant. Assuming *σ* = 0, we can find the lower-bound *α*\* of the mutant apomictic rates such that the mutant invades:

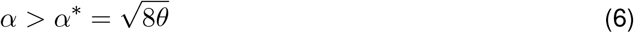

We plot on Fig 5.A *α*\* in function of *c_p_*. As can be seen from Eq 6, *α*\* here does not depend on *c_h_*. This is because in this scenario, apomictic hybrid females can reproduce with both parental and hybrid males while meiotic females cannot reproduce with any of them (see Fig. 3). Therefore, mutant apomictic females gain a reproductive advantage over wild-type meiotic females with independence of the type of mating partner, and thus with independence of hybrid female mating preferences (*c_h_*).

**Figure 5:**
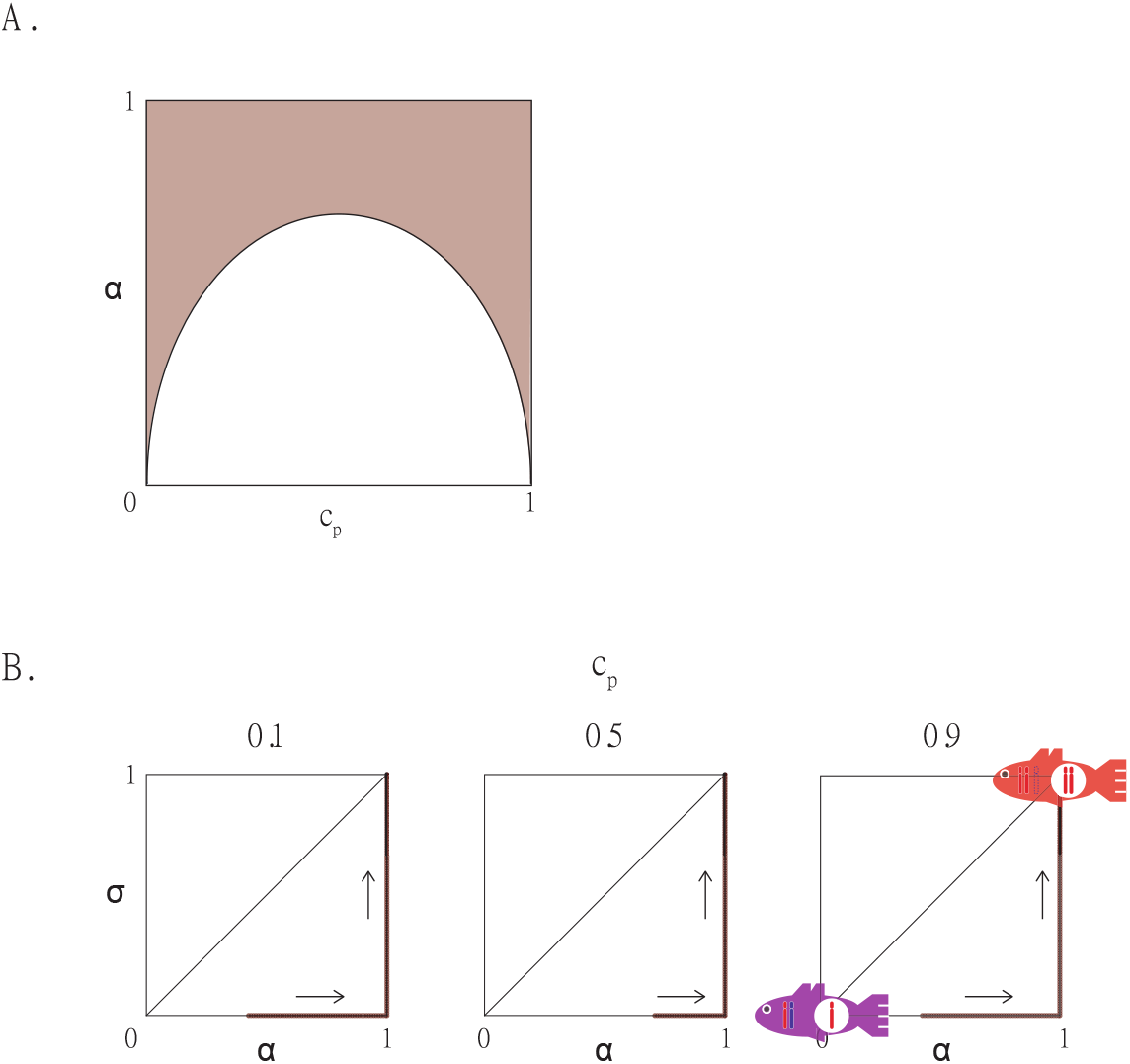
Evolution of gynogenesis in the *egg-fails-to-decondense-sperm scenario*. A. Values of *α* > *α*\* (y-axis) where a mutant can invade are indicated in function of *c_p_* (x-axis) as shaded areas. These do not depend on *c_h_*. B. Predicted evolutionary route to gynogenesis for three values of *c_p_*. This time, *α* appears on the x-axis, while *σ* is represented on the y-axis. These routes show the transition from sexual hybrids (purple fish) to gynogenetic hybrids (red fish). The bottom limit *α*\* of *α* for the first mutant to spread varies in each case, and approximately equal respectively from left to right: (*c_p_* = 0.1) 0.42; (*c_p_* = 0.5) 0.71; (*c_p_* = 0.9) 0.42.

In this scenario, the invasion of a mutant only depends on *θ*, as made clear by Eq 6. As for the previous scenario, *α*\* increases with intermediate values of *c_p_* because these lead to larger values of *θ*, of 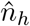, and thus larger mutant death rates by competition for resources with wild-type hybrids.

Once the first mutant with phenotype (*α*\*, *β* = 1, *σ* = 0) has spread and reached its equilibrium population size, we assume that new mutants with any phenotype may appear. We find that the mutant fitness function is again maximized by the gynogenetic phenotype (*α* = 1, *β* = 1, *σ* = 1) (see Supplementary Material for a demonstration).

Noticeably, the partial derivative of the fitness function with respect to *α* is always greater than the partial derivative with respect to *σ* (see Supplementary Material for a demonstration): the selection gradient is always greater in the direction of apomixis than in the direction of female-biased sex-ratios. As such, we expect again in this model evolution to proceed in a stepwise process. Here, apomixis is expected to evolve first, and sex-ratios afterwards. This is illustrated by numerical simulations for different values of *c_p_* shown on Fig 5.B.

### Nonviable-Sperm Scenario

In this scenario, *v* = 1, *s* = 0 and *x* = 1, such that the invasion condition for a mutant female hybrids 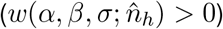 is:

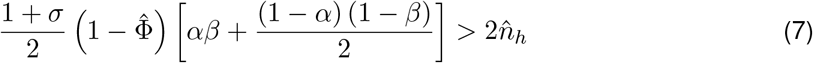

As in the *sperm-fails-to-decondense* scenario, we can use the fact that 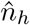 verifies *ṅ_h_* = 0. This time, however, this allows us to show that in this model there can be no invasion of a mutant if *α* = 0, *β* = 0 or *σ* = 0 (see Supplementary Material for a demonstration). Contrarily to the two previous scenarios, here a mutation affecting only apomixis can never spread; it appears that only a mutation affecting the three traits at once can spread. For the mutant to invade, *α* must verify:

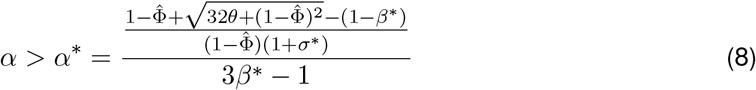

When resolving the inequation *α*\* ≤ 1, we realize that the invasion of a mutant can never happen for *α* ≤ 1/3 and *β* ≤ 1/2. It thus appears that a mutation, in this scenario, must always very significantly alter the reproductive phenotype to be able to invade. This is confirmed by Fig 6.A, where we plot *α*\* as a function of *c_p_* for different values of *c_h_, β* and *σ*: *α*\* is always greater than 1/2, and rapidly increases for decreasing values of *σ* and *β*.

**Figure 6:**
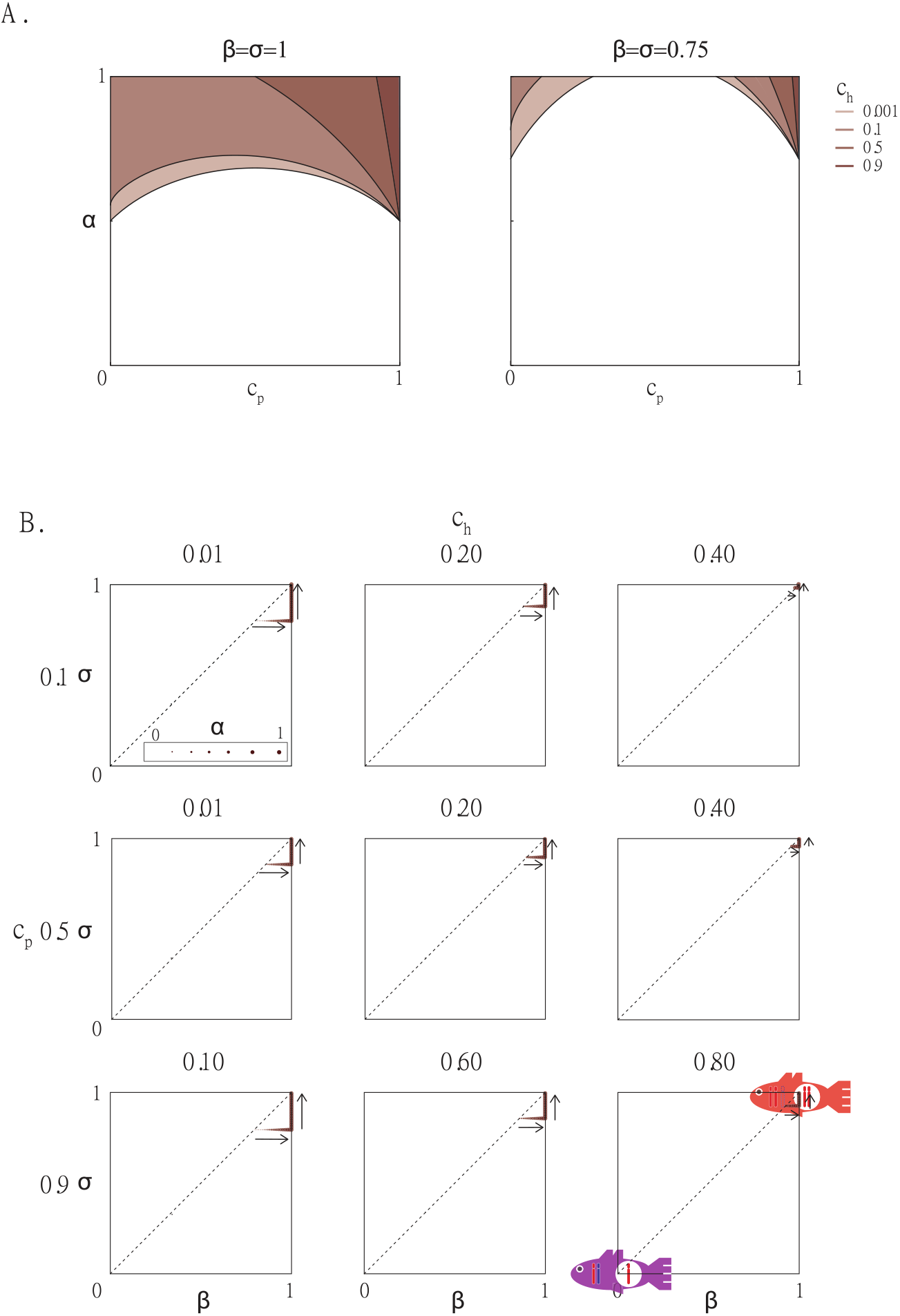
Evolution of gynogenesis in the *nonviable-sperm scenario*. A. Values of *α > α\** (y-axis) where a mutant can invade are indicated in function of *c_p_* (x-axis) and *c_h_* as shaded areas for different values of *β* and *σ*. B. Predicted evolutionary route to gynogenesis for nine combinations of *c_p_* and *c_h_* values. The size of the dot represents *α*, while x- and y-axes represent *β* and *σ* respectively. These routes show the transition from sexual hybrids (purple fish) to gynogenetic hybrids (red fish). The first mutant to spread is assumed to have a mutation such that *α* = *β* = *σ* for simplicity. The bottom limit *α*\*, *β*\* and *σ*\* for the first mutant to spread approximately equals respectively from left to right and top to bottom: (*c_p_* = 0.1) 0.79, 0.88, 0.98; (*c_p_* = 0.5) 0.85, 0.90, 0.96; (*c_p_* = 0.9) 0.80, 0.85, 0.93.

Fig 6.A also shows that again, and for the same reason as before, *α*\* increases with intermediate values of *c_p_*, since, as shown by Eq 8, *α*\* increases with *θ*. Contrarily to the first scenario, *α*\* now increases with *c_h_* (we can see in Eq 8 that *α*\* increases with 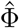). The particularity of this scenario is that a hybrid female, meiotic or apomictic, can never produce viable offspring when mating with a hybrid male. Thus, a mutant can gain a reproductive advantage over wild-types only when mating with parental males, which happens with a probability 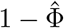 that decreases with *c_h_*. The mutant will gain a reproductive advantage only if it produces apomictic eggs and eliminate paternal genome, which is why the mutation must affect both *α* and *β* at the same time.

Again, the mutant fitness function is maximized by a fully gynogenetic mutant (*α* = 1, *β* = 1, *σ* = 1). Once a first mutant has been able to invade, we thus expect the eventual evolution of gynogenesis by the spreads of new mutants displacing older ones. That is, even though the first step appears difficult in this scenario, if it happens we still predict the eventual spread of gynogenesis via an evolutionary process. We plot on Fig 6.B the expected evolutionary routes to gynogenesis based on fitness gradients for different values of *c_p_* and *c_h_*. We confirm that in this scenario, the first mutant must bring in important alterations of the three traits. Afterwards, we see that apomixis and paternal genome evolution evolve jointly to *α* = 1 and *β* = 1. Once this is reached, sex-ratio evolves in turn to *σ* = 1.

## DISCUSSION

A disproportionate number of transitions from sexuality to asexuality take place in hybrid species [10, 11]. The leading explanation is that genomic incompatibilities between parental genomes result in the spontaneous birth of fully asexual hybrids from sexual parents [13, 14, 15]. This explanation however, is not completely satisfactory. From an empirical perspective, in the rare cases where parental species of hybrid are known and have been crossed, the outcome has been not only asexual but also sexual hybrids. For example, crosses between *Poecilia latipinna* and *Poecilia mexicana*, parental species of the asexual Amazon Molly, have repeatedly failed to produce asexual hybrids [19, 20, 21, 22, 23]. Moreover, genomic signatures suggest that Amazon Mollies crossed with their parental species during their evolutionary history [26] which is incompatible with parental crosses producing asexual hybrids straight away. A similar example in a completely different taxonomic group is the parthenogenetic stick insect *Timema shepardi* [35]. From an evolutionary perspective, the spontaneous generation of asexuals requires genomic incompatibilities that disrupt major reproductive processes without affecting the fertility of hybrids. Here, we explore a more plausible scenario where incompatibilities that disrupt limited parts of the reproductive process lead to the evolution of gynogenesis by natural selection. As such we explore whether natural selection can favour the progressive evolution of asexuals from sexual hybrids with comparatively minor deficiencies in their reproductive system.

Importantly, the previous explanation does not consider that gynogenesis proceeds from sexuality as a result of an evolutionary process. It does not consider whether gynogenesis will be favoured by natural selection against a sexual alternative because this sexual alternative simply does not exist. Our work provides an alternative, evolutionary explanation for the link between hybridization and asexuality. By formally considering natural selection between sexual and asexual hybrids, we provide a more satisfactory theory from an evolutionary perspective.

Our work allows for the different reproductive phenotypes that characterize gynogenesis to evolve independently. We decouple the two components of amphimixis, that is, the fusion of the egg and sperm that triggers embryogenesis from the union of maternal and paternal genomes that traditionally restores ploidy. The three models we have analyzed represent two very different forms of fertilization disruption as a result of hybridization. In the first two scenarios, hybrid egg and sperm can fuse and embryogenesis can be triggered, but the union of maternal and paternal genome fails. In the first model, it fails because hybrid sperm is unable to provide the necessary machinery (for example, it is unable to do its part in the decondensation of the sperm’s pronucleus). In the second scenario, it fails because hybrid eggs are unable to provide the necessary machinery (again, for example, they are unable to participate to sperm pronucleus decondensation). In contrast, in the third scenario, hybrid egg and sperm cannot even fuse, and thus hybrid embryogenesis cannot be triggered by hybrid sperm, for example because hybrid males are unable to produce viable sperm. We show how gynogenesis can evolve from any of these disruptions. We thus propose a new link between hybridization and asexuality: that hybridization disrupts one of the two aspects of amphimixis, which in turns triggers the evolution of asexuality as a way to rescue hybrid reproduction. Importantly, our models show that maternal and paternal genome union failure (whether it originates from the egg or from the sperm) is a more favourable scenario for the evolution of asexuality, especially considering that it allows for the independent evolution of unreduced meiosis and paternal genome elimination. In contrast, egg and sperm fusion failure – that is, failure to even enter embryogenesis – requires a potentially unlikely coincidence of events (large mutations affecting the three phenotypic traits at once) for gynogenesis to evolve. Whichever step is disrupted, such disruptions are arguably much more susceptible to appear in hybrid species that combine diverged parental genomes than in non-hybrid species, which would explain why so many asexual species are of hybrid origin. That is, we propose that many asexual species are hybrid species because hybrid species are a fertile ground for the evolution of asexuality by natural selection: hybrid species have more chances to have limited reproductive capabilities, which may be rescued by the evolution of asexuality in certain cases.

To the best of our knowledge, there exists no conclusive evidence that shows how failure in the union of maternal and paternal genomes may happen in hybrid species (including crosses between *P. mexicana* and *P. latipinna*). This is because it is a complex cytological mechanism and until now there was no reason to specifically look for the mechanics of such disruption in hybrids; we hope our work will raise a stronger interest in investigating this. Although there are no specific examples of union failure in hybrids due to paternal or maternal factors, there are various indirect evidence that support our hypothesis. On one hand, a diversity of cases where progeny of interspecific crosses did not incorporate paternal genetic material have been recorded [36, 37, 38]. On the other hand, there is evidence that egg products can prevent sperm pronucleus decondensation. For example, sperm pronuclei of the sexual Common Carp (*Cyprinus carpio*) is unable to decondense in egg extracts of the gynogenetic Gibel Carp (*Carassius auratus gibelio*) [39]. Also, there is evidence that changes in the paternal genome can be responsible for maternal and paternal genome union failure. For example, non-functional mutations in Drosophila males (*Drosophila melanogaster*) lead to sperm pronucleus decondensation failure [40, 41]. Overall, several lines of evidence support the plausibility of the assumptions underlying the scenarios that involve a failure of the union of maternal and paternal genomes. The assumption of the third model, that hybrid sperm is nonviable, has been frequently observed and relates to Haldane’s rule [42, 43].

Our work shows that natural selection may favour the progressive (one phenotypic trait at a time) evolution of gynogenesis. We predict that when hybrid sperm can trigger embryogenesis but fails to decondense its own pronucleus after fusing with an egg (first scenario), apomixis can evolve independently of other traits because apomictic females can produce viable diploid progeny when mating with hybrid males even without eliminating paternal genomes. We show that production of more females than males follows. Paternal genome elimination is not required for any of these features to evolve and can evolve subsequently to allow apomictic females to produce viable offspring when mating with parental males too, which becomes increasingly necessary as hybrid sex-ratio gets more and more female-biased. Similarly, we predict that when hybrid sperm can trigger embryogenesis but hybrid eggs fail to decondense sperm pronuclei (second scenario), apomixis evolves independently because it restores hybrid female fertility. A female bias sex-ratio evolves afterwards. Paternal genome elimination does not evolve as the paternal genome is never transmitted by hybrid females anyway. Finally, we predict that when hybrids produce nonviable sperm (third scenario), apomixis cannot evolve independently. Evolution only favours the spread of the three traits together, simultaneously. In this model the conditions for gynogenesis to evolve are very restrictive. In particular, a fully, or almost fully gynogenetic hybrid has to appear as a result of a mutation, which has been claimed to be difficult [44]. Importantly, in the first two scenarios, we propose for the first time an evolutionary path where unreduced meiosis and paternal genome elimination does not necessarily need to evolve together, which significantly simplify the underlying evolutionary processes. As such, we expect these scenarios to be particularly prone to the evolution of gynogenesis in particular and asexuality in general (see below).

The selective force driving the evolution of gynogenesis is a form of reproductive assurance. *Reproductive assurance theory* contends that true parthenogenetic females gain a reproductive advantage over sexual ones because they do not need mating and fertilization (which may fail for a diversity of reasons) [45]. Here, we show that this idea of reproductive assurance can also apply to the evolution of gynogenesis. Even though gynogens do need mates and fertilization (egg and sperm fusion), we show that they can still benefit from reproductive assurance if male gametes are not always able to transmit their genomes (in which case sexual females have limited reproductive outputs). Traditionally, reproductive assurance has been associated with mate scarcity because of reduced population sizes or biased sex-ratios [46, 47, 48, 49]. Here, we argue that mates may not be lacking, but mates able to transfer their genetic material may be because of disruptions to the union of maternal and paternal genomes. Whenever this happens, we expect gynogenesis to receive a selective advantage.

Our research underscores the importance of the mating structure on the evolution of gynogenesis. We showed that random mating between parental species, that is intermediate values of *c_p_*, makes the evolution of gynogenesis most difficult in the three scenarios (see Fig 4, 5 and 6). However, we emphasized that this effect is really about asexual mutants having a harder time invading whenever there is a greater production of sexual hybrids by direct hybridization between the parental species. Thus, we predict that the evolution of asexuality in hybrids should rather be expected in cases of rare hybridization, whether it is because of mating choices (for example, large *c_p_*), or because of the geographical distributions of the parental species. Interestingly, modelling work has recently suggested that there has been little historical overlap between the geographical distributions of *P. mexicana* and *P. latipinna* [50], which points towards historically rare hybridization events among the ancestors of the Amazon Molly. In the *sperm-fails-to-decondense scenario*, assortative mating within hybrids, that is *c_h_* ≈ 1, most favours the evolution of gynogenesis (see Fig 4) because male gametes unable to transmit their genomes allow apomictic females to gain a reproductive advantage over meiotic females. In the *egg-fails-to-decondense-sperm scenario*, the likeliness that gynogenesis will evolve does not depend on hybrid matings (see Fig. 5), because apomixis provides a reproductive advantage independently of with whom the females are mating. Finally, in the *nonviable-sperm scenario*, gynogenesis is most likely to evolve in cases of disassortative mating within hybrids (*c_h_* ≈ 0, see Fig 6), because hybrid males cannot trigger embryogenesis of meiotic nor apomictic females.

We would like to conclude by emphasizing that though we focused here on gynogenesis, the model we built can actually be extended to the evolution of true (thelytokous) parthenogenesis. Parthenogenetic species do not perform paternal genome elimination, but instead benefit from spontaneous embryogenesis: they do not need the fusion of egg and sperm for embryogenesis to start (and thus, they do not need to eliminate any paternal genome). Mathematically, spontaneous embryogenesis can be treated exactly as paternal genome elimination in our model. For example, in our model, a meiotic female can reproduce with a parental male if it does not eliminate paternal genome, and an apomictic female can reproduce with a parental male if it does eliminate paternal genome (see Fig 3). This can perfectly be translated into parthenogenetic traits. On one hand, a meiotic female can reproduce with a parental male only if its egg does not spontaneously enter embryogenesis – in which case the sperm would not have had time to fuse with the egg and contribute its genome; the offspring would be haploid and unviable. On the other hand, an apomictic female can reproduce with parental males only if its egg does spontaneously enter embryogenesis before any fertilization has had a chance to take place – in which case the sperm would fuse with the egg, contribute its genome, resulting in the production of a triploid, unviable offspring. Because paternal genome elimination and spontaneous embryogenesis can be seen as two ways of doing the same thing – beginning embryonic divisions without the fusion of maternal and paternal genomes – we argue that our findings apply not only to the evolution of gynogenesis, but also to the evolution of true parthenogenesis. Overall, we proposed here a novel theory for the evolution of asexuality in general, providing for the first time an evolutionary link between asexuality and hybridization, and showing that in certain cases unreduced meiosis can evolve independently of paternal genome elimination (or spontaneous embryogenesis), significantly easing the evolutionary process.

