## Supplementary Material for "Why Do Hybrids Turn Down Sex?"

#### Contents

**1 Full description of the model** **2**

**2 Equilibrium size of the parental populations** **4**

**3 Equilibrium size of the wild-type hybrid population** **5**

**4 Scenario 1 - Sperm fails to decondense** **7**

4.1 Invasion condition of a partially asexual mutant . . . . . . . . . . . . . . . . . . . . . . 7

4.2 Wild-Type - Mutant hybrid equilibrium . . . . . . . . . . . . . . . . . . . . . . . . . . 8

4.3 Evolutionary route to asexuality . . . . . . . . . . . . . . . . . . . . . . . . . . . . . . 9

**5 Scenario 2 - Egg fails to decondense the sperm** **11**

5.1 Invasion condition of a partially asexual mutant . . . . . . . . . . . . . . . . . . . . . . 12

5.2 Wild-Type - Mutant hybrid equilibrium . . . . . . . . . . . . . . . . . . . . . . . . . . 12

5.3 Evolutionary route to asexuality . . . . . . . . . . . . . . . . . . . . . . . . . . . . . . 12

**6 Scenario 3 - Nonviable sperm** **14**

6.1 Invasion condition of a partially asexual mutant . . . . . . . . . . . . . . . . . . . . . . 14

6.2 Wild-Type - Mutant hybrid equilibrium . . . . . . . . . . . . . . . . . . . . . . . . . . 15

6.3 Evolutionary route to asexuality . . . . . . . . . . . . . . . . . . . . . . . . . . . . . . 15

**7 Relaxing the absence of inter-specific competition** **15**

**8 Relaxing the complete sexuality of wild-type hybrids** **18**

### FULL DESCRIPTION OF THE MODEL

We consider three populations coexisting in a common environment: two parental species $(1)$ and $(2)$, along with a hybrid population that results from the interbreeding of the two parental species $(h)$.

We assume that the three species coexist in a same environment and potentially compete for resources. We also assume that the parental populations only coexist in a margin of their respective ecological distribution. As a result, parental populations' global dynamics should be negligibly influenced by competition between each other and with the hybrid species. That is, the death rate $\Psi_i$ of any population $i \in \{1, 2\}$ is only a function of intra-specific competition for resources:

$$\Psi_i = \frac{N_i^{♀} + N_i^{♂}}{K_i} \tag{1}$$

with $N_i^{♀}$ the number of females from population $i$ in the environment, $N_i^{♂}$ the number of males from population $i$ in the environment, and $K_i$ the carrying capacity of the environment for population $i$.

The whole hybrid species, in contrast, coexist with parental competitors over all its ecological distribution. We note $\chi$ the parameter that reflects the intensity of this inter-specific competition. For hybrids, equation 1 becomes:

$$\Psi_h = \frac{N_h^{♀} + N_h^{♂}}{K_h} + \chi \sum_{j \in \{1,2\}} \frac{N_j^{♀} + N_j^{♂}}{K_j} \tag{2}$$

We show later that the competition parameter $\chi$ does not play a critical role in the outcome of the model. Competition for resources from parentals to hybrids may lead to the extinction of the hybrid species. Here we are not interested in the conditions that ensure or not the coexistence of the hybrid and parental populations. In the following, we will assume that $\chi = 0$ most of the time, as it is a simple way to ensure coexistence that does not appear to qualitatively alter our results.

Because we assume that the parental species and the wild-type hybrids are completely sexual and exhibit unbiased sex-ratios, we note in the following $N_i = N_i^{♀} = N_i^{♂}$. As such, $N_i$ really represents half of the total size of population $i$. Also, we consider that the parental populations have the same behaviour; as such, we assume that they have equal population sizes at all time: $N_1 = N_2$. We can group them under the subscript $p$: $N_p = N_1 = N_2$.

Henceforth, we assume for simplicity that $K_1 = K_2 = K_h = K$. To further simplify the equations, we change variables and call $n_i$ the size of population $i \in \{1, 2, h\}$ relative to the carrying capacity of the environment: $n_i = N_i/K$. Overall, we thus have $\Psi_i = 2n_i$.

Each population is characterized by certain mating preferences: we note $c_{ij}$ the relative preference of females from population $i$ to males from population $j$. High values of $c_{ii}$ correspond to assortative mating patterns: a female from a given population prefers to mate with males from the same population over males from the other populations. Intermediary $c_{ii}$ relates to random mating, while low values of $c_{ii}$ represent disassortative mating, a scenario where females actually prefer to mate with males from other populations. We note $\Phi_{ij}$ the probability of females from population $i$ to mate with males of population $j$:

$$\Phi_{ij} = \frac{c_{ij} n_j}{\sum_k c_{ik} n_k} \tag{3}$$

$c_{ij}$ are defined such that $\forall i \in \{1, 2, h\},\ \sum_k c_{ik} = 1$.

We make some simplifying assumptions regarding the mating choices to reduce the number of parameters. First, we consider that the two parent species exhibit similar levels of assortative mating: $c_{11} = c_{22} = c_p$. Also, we assume that parent females never try to mate with hybrid males, as such matings do not bear viable offspring. That is, $c_{1h} = c_{2h} = 0$. Thus $c_{12} = c_{21} = 1 - c_p$. Concerning the hybrid species, we note $c_{hh} = c_h$ and assume that hybrid females do not discriminate between the parent species, such that $c_{h1} = c_{h2} = (1 - c_h)/2$.

As a result from this assumption, $\Phi_{h1} = \Phi_{h2} = \dfrac{1 - \Phi_{hh}}{2}$. In the following and in the main text, we call simply $\Phi$ the probability of a hybrid female mating with a hybrid male ($\Phi_{hh}$). Matings of hybrid females with males from any of the two parental populations are grouped together and occur with probability $1 - \Phi$. Following these assumptions, we have:

$$\Phi = \frac{c_h n_h}{c_h n_h + (1 - c_h) n_p} \tag{4}$$

### EQUILIBRIUM SIZE OF THE PARENTAL POPULATIONS

The parental system's dynamics can be described using the following Ordinary Differential Equations (ODEs):

$$\frac{\mathrm{d}n_p}{\mathrm{d}t} = r_p \left( \frac{1}{2} \frac{c_p n_p}{c_p n_p + (1 - c_p) n_p} - 2n_p \right) n_p = r_p \left( \frac{c_p}{2} - 2n_p \right) n_p \tag{5}$$

with $r_p$ referring to the number of progeny obtained by parental females at each generation. In the following, we will assume for simplicity that all populations have the same intrinsic growth rate: $r_p = r_h = r$.

Solving this ODE, we can obtain the equilibrium of this parental system such that the two species coexist:

$$\hat{n}_p = \frac{c_p}{4} \tag{6}$$

We can calculate the partial derivative of Eq. (5) with respect to $n_p$ at equilibrium:

$$\left. \frac{\partial \dot{n}_p}{\partial n_p} \right)_{n_p = \hat{n}_p} = \frac{c_p}{2} - 4\hat{n}_p = -\frac{c_p}{2} < 0 \tag{7}$$

with $\dot{n}_p = \dfrac{\mathrm{d}n_p}{\mathrm{d}\tau}$ and $\tau = tr$.

The partial derivative being negative in the vicinity of the equilibrium, we can conclude that this equilibrium is asymptotically stable.

**As a simple system of birth and death by competition for resources, the parental populations reach a stable equilibrium population size of $\hat{n}_p = \frac{c_p}{4}$. Throughout the rest of the analysis, we will simply assume that parental species always are at this equilibrium.**

### EQUILIBRIUM SIZE OF THE WILD-TYPE HYBRID POPULATION

We consider two possible hybrid populations. The wild-type hybrid population is a completely sexual population. The mutant population, however, may exhibit asexual traits, namely the production of clonal eggs with probability $\alpha$ ($\alpha \in [0,1]$), spontaneous embryogenesis / paternal genome elimination with probability $\beta$ ($\beta \in [0,1]$) and the production of progeny with sex-ratio $\frac{1+\sigma}{2}$ ($\sigma \in [-1,1]$).

Wild-type hybrids can be born by (1) direct hybridization between the parental species and (2) back-crossing of meiotic hybrid females (wild-type or mutant) with parental species. Wild-type females produce exclusively wild-type hybrids when backcrossing. Backcrossing mutants are assumed to produce half of sexual wild-type hybrids and half of partially asexual mutant hybrids (because they mated with sexual parental males). The mating structure and viability of progeny depends on the scenario we consider, and are detailed in the main text.

Hybridization happens with probability $(1-c_p)n_p = \frac{c_p(1-c_p)}{4}$. We note $\theta$ this constant flow of wild-type hybrids due to direct hybridization between the parental populations. Assuming that at first there is no mutant hybrid yet, the dynamics of the wild-type hybrid population can be described as:

$$\dot{n}_h = \theta + \left(\frac{1-\Phi}{2} - 2n_h\right) n_h \tag{8}$$

in *sperm fails to decondense* and *nonviable sperm* scenarios. Here, $\dot{n}_h$ refers to $\frac{\mathrm{d}n_h}{\mathrm{d}\tau}$. In contrast, in the *egg fails to decondense sperm* scenario, backcrossings of wild-type females with parental males are unproductive, and the ODE becomes:

$$\dot{n}_h = \theta - 2n_h^2 \tag{9}$$

We can write a general equation that works for the three scenarios:

$$\dot{n}_h = \theta + \left(v\frac{1-\Phi}{2} - 2n_h\right) n_h \tag{10}$$

with $v = 0$ for the *egg fails to decondense sperm* scenario, and $v = 1$ for the other two scenarios.

In *sperm fails to decondense* and *nonviable sperm* scenarios, we cannot find an analytical equation for the equilibrium population size of wild-type hybrids $\hat{n}_h$. We can however find an expression of it that depends on $\hat{\Phi}$ (which itself depends on $\hat{n}_h$, so this is not a closed-form expression):

$$\hat{n}_h = \frac{1 - \hat{\Phi} + \sqrt{32\theta + \left(1 - \hat{\Phi}\right)^2}}{8} \tag{11}$$

In the *egg fails to decondense sperm* scenario, however, a closed-form expression of $\hat{n}_h$ can easily be found:

$$\hat{n}_h = \sqrt{\frac{\theta}{2}} \tag{12}$$

The partial derivative of $\dot{n}_h$ with respect to $n_h$ at equilibrium is:

$$\left.\frac{\partial \dot{n}_h}{\partial n_h}\right)_{n_h=\hat{n}_h} = v\frac{1-\hat{\Phi}}{2} - 2\hat{n}_h - n_h\left(\frac{v}{2}\left.\frac{\partial \dot{\Phi}}{\partial n_h}\right)_{n_h=\hat{n}_h} + 2\right) \tag{13}$$

$\frac{\partial \dot{\Phi}}{\partial n_h}$ can easily be shown to be always positive: as there are more hybrids, hybrids $\times$ hybrids matings are more probable. In addition, $v\frac{1-\Phi}{2} - 2\hat{n}_h \leq 0$ by definition of $\widehat{n_h}$ ($\dot{n}_h = 0$). Thus, the equilibrium found here is stable.

> **The wild-type hybrid population reaches a stable equilibrium size $\hat{n}_h$ in the three scenarios. It is not always possible to obtain a closed-form expression of this equilibrium.**

### SCENARIO 1 - SPERM FAILS TO DECONDENSE

In this model, we assume that all hybrid males have functional sperm and that spermatozoa are able to bind and fuse with oocytes from all types of females – such that they trigger embryogenesis. However, we also assume that the hybrid males' sperm is sterile in the sense that spermatozoa are unable to decondense their pronuclei and transmit to the eggs their genetic material. As a result, if the egg is haploid, the resulting embryo will still be haploid and unviable. On the contrary, if the egg is diploid, the embryo will be diploid and considered viable.

#### 4.1 Invasion condition of a partially asexual mutant

Once the wild-type hybrids have reached their equilibrium, we introduce a rare (such that $n_h = \hat{n}_h$ is still approximately true) mutant hybrid in the environment. This mutant population has a phenotype $\{\alpha, \beta, \sigma\}$. The growth rate – in the following, we call this fitness – of such a mutant population is:

$$w = \frac{1+\sigma}{2}\left[\alpha\left(\hat{\Phi} + \left(1-\hat{\Phi}\right)\beta\right) + (1-\alpha)\frac{1}{2}\left(1-\hat{\Phi}\right)(1-\beta)\right] - 2\hat{n}_h \tag{14}$$

The invasion of a mutant hybrid is possible if and only if $w > 0$. This translates into:

$$\frac{1+\sigma}{2}\left(\hat{\Phi} + \frac{(1-\hat{\Phi})(\beta - 1/3)}{2/3}\right)\alpha > 2\hat{n}_h - \frac{(1+\sigma)(1-\beta)}{2}\frac{1-\hat{\Phi}}{2} \tag{15}$$

Now, notice that the right-hand side of this condition is greater than $2\hat{n}_h - \frac{1-\hat{\Phi}}{2}$, which we know is positive (see above). Thus, the left-hand side of Eq. (15) must necessarily be strictly positive for the invasion condition to potentially be met. This in turn necessarily means that $\alpha$ must be greater than $0$ for a mutant to have a chance to invade.

Assuming that any mutation can only alter one phenotypic trait, we thus understand that the first mutation to invade must alter $\alpha$. Posing $\beta = \sigma = 0$, we find that Eq. (15) can be rewritten as:

$$\alpha > \alpha^* = \frac{8\hat{n}_h - (1-\hat{\Phi})}{3\Phi - 1} \tag{16}$$

or, substituting $\hat{n}_h$ by its expression in function of $\Phi$:

$$\alpha > \alpha^* = \frac{\sqrt{32\theta + (1 - \hat{\Phi})^2}}{3\hat{\Phi} - 1} \tag{17}$$

There is a minimal value of $\alpha$ under which a mutant cannot spread. However, this value is not necessarily within $[0, 1]$. $\alpha^* < 0$ if $\hat{\Phi} < 1/3$. However, when $\hat{\Phi} < 1/3$, the left-hand side of Eq. (15) cannot be positive; there can never be the invasion of a mutant hybrid if $\hat{\Phi} < 1/3$. We can thus focus on $\hat{\Phi} > 1/3$. If $\Phi$ is close to $1/3$ or $\theta$ large, then it is possible that $\alpha^* > 1$. In that case, no mutant can possibly invade. As such, invasion is rather possible when $\hat{\Phi}$ is large ($c_h$ large) and $\theta$ is small (extreme values of $c_p$).

**Any mutant must have a strictly positive rate of clonal egg production to potentially invade. Such mutant tends to invade when $c_h$ is large and $c_p$ extreme (close to $0$ or $1$).**

#### 4.2 Wild-Type - Mutant hybrid equilibrium

When a mutant invade, it reaches a polymorphic equilibrium where both wild-type and mutant hybrids coexist. Wild-type are not completely eliminated from the environment because they are still produced by direct hybridization and back-crossing of mutant females with parental males. We can show numerically that this equilibrium is stable. We provide below a few examples of mutant invasion.

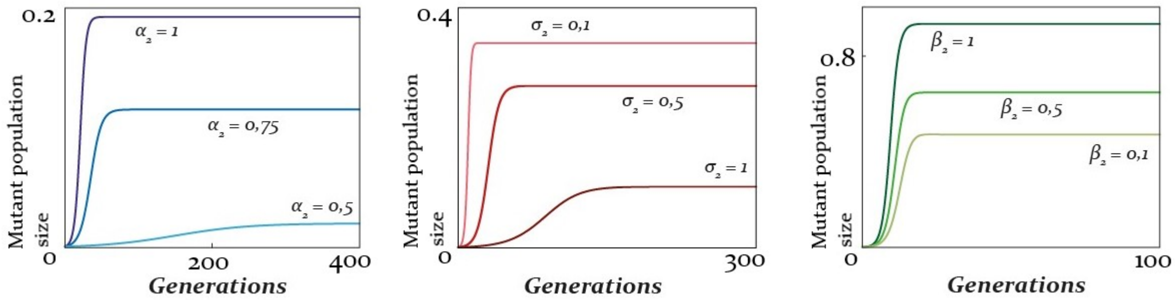


**Figure 1:** Number of mutant hybrid females through time, depending on different parameter values. Initial conditions are: wild-type hybrids at equilibrium, mutant hybrids with only $1$ male and $1$ female. *(a)* $\beta = 0$, $\sigma = 0$, $c_p = 0.1$ and $c_h = 0.8$. This case corresponds to the first partially apomictic mutant (with $\alpha^* \approx 0.457$ for these parameter values). *(b)* $\alpha = 1$, $\beta = 0$, $c_p = 0.9$ and $c_h = 0.85$.*(c)* $\alpha = 1$, $\sigma = 1$, $c_p = 0.1$ and $c_h = 0.5$. The value of the varying trait is indicated in each graph, with darker color indicating higher trait values.

#### 4.3 Evolutionary route to asexuality

Once a mutant has been able to spread, any alternative mutant that has a reproductive benefits over it will in turn spread and displace it. This leads us to realize that derivative of $w$ with respect to the phenotypic trait of the new mutant $\alpha'$, $\beta'$ and $\sigma'$ actually gives us the selection gradient applying to mutant hybrids. Note that here $\hat{\Phi}$ and $\hat{n}_h$ now refers to the wild-type - mutant equilibrium, which depends on $c_h$, $c_p$ and the phenotypic traits of the established mutant $\alpha$, $\beta$ and $\sigma$.

These partial derivatives are:

$$\begin{cases} \dfrac{\partial w}{\partial \alpha'} = \dfrac{(1+\sigma)\left(3\hat{\Phi} - 1 + 3\beta(1-\hat{\Phi})\right)}{4} & \text{(18a)} \\ \dfrac{\partial w}{\partial \beta'} = \dfrac{(1+\sigma)(1-\hat{\Phi})(3\alpha - 1)}{4} & \text{(18b)} \\ \dfrac{\partial w}{\partial \sigma'} = \dfrac{(1-\alpha)(1-\beta)(1-\hat{\Phi}) + 2\alpha(\hat{\Phi} + \beta(1-\hat{\Phi}))}{4} & \text{(18c)} \end{cases}$$

The sign of Eq (18a) only depends on the sign of $3\hat{\Phi} - 1 + 3\beta(1-\hat{\Phi})$. This is positive if either $\hat{\Phi}$ or $\beta$ are large enough (worst-case scenario, one or the other larger than $1/3$). This is because there are two ways that clonal eggs can produce enough viable diploid offspring: either because eggs often do not incorporate sperm pronuclei ($\beta$ large enough) or because females often mate with hybrid males unable to transmit their genome ($\hat{\Phi}$ large enough). $3\hat{\Phi} - 1 + 3\beta(1-\hat{\Phi})$ cannot be negative if $\hat{\Phi} > 1/3$. We can intuitively understand, without deriving expression for $\hat{n}_h$ and $\hat{\Phi}$ at the wild-type - mutant equilibrium, that the spread of mutant hybrids tends to increase the overall amount of hybrids since the mutation brings in additional reproductive capabilities (if not, it could not invade). Thus, we know that $\hat{n}_h$ is greater at the wild-type mutant equilibrium than at the wild-type only equilibrium. As such, $\hat{\Phi}$ must also be greater at the wild-type equilibrium than at the wild-type only equilibrium. Because we had $\hat{\Phi} > 1/3$ at the wild-type equilibrium for a mutant to potentially invade, we know that at the wild-type - mutant equilibrium we necessarily also have $\hat{\Phi} > 1/3$. Thus, Eq. (18a) is always positive: mutant fitness is maximized for $\alpha' = 1$; females producing exclusively clonal eggs should eventually invade.

It is trivial to see that Eq. (18c) is always positive : $w$ is maximized at $\sigma' = 1$, such that mutant hybrid females exclusively bearing females should eventually spread. This is straightforward because males are sterile and do not transmit their genes, such that it's always beneficial in this system to produce more females and less males. Finally, Eq. (18b) is positive if and only if $\alpha > 1/3$. Because we saw above that $100\%$ clonal egg production is eventually expected to evolve once a first partially clonal mutant has spread, then $\alpha$ will ultimately be greater than $1/3$, and thus Eq. (18b) will eventually be positive. $w$ is maximized at $\beta = 1$: systematic paternal genome elimination is expected to spread.

**If a first partially clonal mutant can spread, selection will favour the spread of new mutations (which can be arbitrarily small) that will eventually lead to the rise and spread of a fully asexual class of hybrids ($\{\alpha = 1, \beta = 1, \sigma = 1\}$). Spontaneous embryogenesis / paternal genome elimination can spread only once rates of clonal egg production have reached a minimal value of $1/3$.**

The evolutionary dynamics of this system is characterized by the recurrent invasion of new mutants that spread, reach a stable population size, before being in turn replaced until a complete asexual mutant has invaded. We show on Fig. 2 a numerical example of this mutation and replacement process. As more asexual new mutants spread, the overall number of mutant hybrids increase, while the number of wild-type hybrids decreases. Once full asexuality has spread, the wild-type hybrids are only maintained by direct hybridization between the parental species, and are thus expected to ultimately disappear once these have diverged so much that they cannot hybridize anymore.

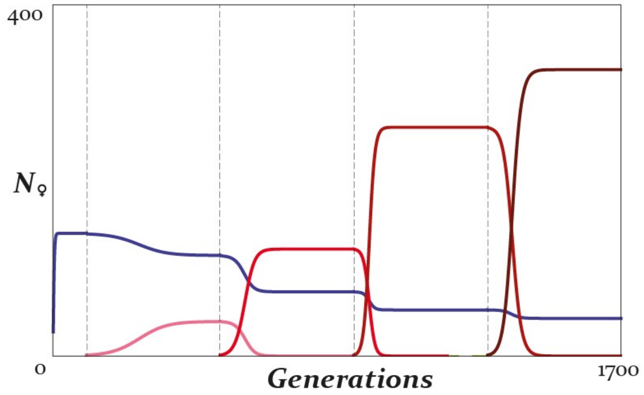


**Figure 2:** Numerical simulations of wild-type (blue) and mutant (red) hybrid populations. Initially, only the parent species are present. At $t = 1$, one wild-type hybrid male and one wild-type hybrid female are introduced. The wild-type population increases logistically until reaching an equilibrium. At $t = 100$, we introduce one mutant male and one mutant female. This first mutant is partially clonal, characterized by $\{\alpha_2 = 0.8, \beta_2 = 0, \sigma_2 = 0\}$. The mutant population increases until reaching an equilibrium - in parallel, the wild-type hybrid population decreases in size until reaching an equilibrium. Every $400$ generations, a new more asexual mutant is introduced (darker shades of red). $t = 500$: $\{\alpha_3 = 1, \beta_3 = 0, \sigma_3 = 0\}$; $t = 900$: $\{\alpha_3 = 1, \beta_3 = 0, \sigma_3 = 0.3\}$; $t = 1300$: $\{\alpha_3 = 1, \beta_3 = 0.6, \sigma_3 = 0.3\}$. Each time the new mutant population size increases logistically, reaches an equilibrium population size, and drives older, less asexual mutants to extinction. The invasion of each new mutant further decreases wild-type hybrid population size.

### SCENARIO 2 - EGG FAILS TO DECONDENSE THE SPERM

In this model, we assume hybrid males also have functional sperm and spermatozoa are still able to bind and fuse with oocytes from all types of females to trigger embryogenesis. However, this time we assume that hybrid eggs are unable to decondense and incorporate the genetic material of any kind of sperm. As a result, as in the previous scenario, if the egg is haploid, the resulting embryo will still be haploid and unviable. On the contrary, if the egg is diploid, the embryo will be diploid and considered viable.

In the following, we follow the same steps as for the previous scenario. As we have already gone over the whole process for the previous scenario, in the following we omit to define the variables. we ask the reader to refer to the previous section for definitions of the variables and more ample justification of the method we follow.

#### 5.1 Invasion condition of a partially asexual mutant

In this model, the fitness of a mutant can be written as:

$$w = \frac{1+\sigma}{2}\alpha - 2\hat{n}_h \tag{19}$$

It is straightforward to see that $w > 0$ again implies that $\alpha > 0$: a mutation must necessarily involve the production of clonal eggs to have a chance to invade. Assuming that such a mutant only involves the production of clonal eggs ($\sigma = 0$), the lower-bound of the mutation effect for the mutant to invade is:

$$\alpha > \alpha^* = 4\hat{n}_h = \sqrt{8\theta} = \sqrt{2c_p(1-c_p)} \tag{20}$$

Note that in this scenario, $0 \geq \alpha^* \geq 1$ for all values of $c_p$ and $c_h$.

**Any mutant must have a strictly positive rate of clonal egg production to potentially invade. For all parameter values, there can be the invasion of some mutant. The size of the necessary mutation for invasion is greater at intermediate values of $c_p$ (more influx from hybridization), and does not depend on $c_h$.**

#### 5.2 Wild-Type - Mutant hybrid equilibrium

Again, we can show both numerically and analytically that if the mutant can spread, it will reach a stable equilibrium population size.

#### 5.3 Evolutionary route to asexuality

The partial derivatives of the fitness function in this model are:

$$\begin{cases} \dfrac{\partial w}{\partial \alpha'} = \dfrac{1+\sigma}{2} \geq 0 & \text{(21a)} \\ \dfrac{\partial w}{\partial \sigma'} = \dfrac{\alpha}{2} \geq 0 & \text{(21b)} \end{cases}$$

Eq. (21a) and (21b) are always trivially positive. It follows that the fitness function is maximized by a fully asexual mutant: $\alpha = 1$ and $\sigma = 1$ ($\beta = 1$ for all hybrids by construction of this model).

Interestingly, $\alpha$ is by definition always smaller than $1 + \sigma$ provided that $\sigma > 0$ ($\sigma < 0$ is not expected since the partial derivative of the fitness function with respect to $\sigma'$ is always positive). Thus, $\frac{\partial w}{\partial \alpha'}$ is always greater than $\frac{\partial w}{\partial \sigma'}$. In other words, the selection force on clonal egg production is expected to be always greater than the one affecting sex-ratios. As a result, we expect in general clonal egg production to fully evolve before female-biased sex-ratios do.

**Once a first partially clonal mutant has spread, we expect evolution to select for increasingly asexual mutants, until full asexuality is reached. In this scenario, spontaneous embryogenesis / paternal genome elimination appears as a result of the genetic incompatibilities between the parental species. If the necessary conditions are met (a mutation whose effect is greater than $\sqrt{8\theta}$), then clonal egg production fully evolve first, and female-biased sex-ratios second.**

### SCENARIO 3 - NONVIABLE SPERM

In this model, hybrid males are unable to produce viable sperm and, as such, are unable to trigger embryogenesis. As a result, hybrid female eggs – haploid or diploid – can only start embryogenesis when the females mate with parental males.

Again, we refer the reader to section 1 for ample description of the variables and the method followed.

#### 6.1 Invasion condition of a partially asexual mutant

In this model, the fitness of a mutant can be written as:

$$w = \frac{1+\sigma}{2}\left[\alpha\left(1-\hat{\Phi}\right)\beta + (1-\alpha)\frac{1}{2}\left(1-\hat{\Phi}\right)(1-\beta)\right] - 2\hat{n}_h \tag{22}$$

$w > 0$ now translates into:

$$\frac{1+\sigma}{2}\alpha\frac{3\beta-1}{2}\left(1-\hat{\Phi}\right) > 2\hat{n}_h - \frac{1}{2}\left(1-\hat{\Phi}\right)(1-\beta) \tag{23}$$

We can use $\dot{n}_h = 0$ to show that at equilibrium the right-hand side of Eq. (23) is greater that $\frac{1-\hat{\Phi}}{2}$. A mutant must thus verify:

$$\frac{1+\sigma}{2}\alpha(3\beta-1) > 0 \tag{24}$$

Note that this can happen if and only if $\alpha > 0$ and $\beta > \frac{1}{3}$. The invasion condition can be written in terms of $\alpha^*$, $\beta^*$ and $\sigma^*$:

$$\alpha > \alpha^* = \frac{\frac{1-\hat{\Phi}+\sqrt{32\theta+(1-\hat{\Phi})^2}-(1-\beta^*)}{(1-\hat{\Phi})(1+\sigma^*)}}{3\beta^*-1} \tag{25}$$

A sign analysis reveals that $\alpha^* \geq 1$ if and only if $\beta^* \geq \frac{1}{2}$.

**Any mutant must affect both $\alpha$ and $\beta$ at once to be able to invade.**

#### 6.2 Wild-Type - Mutant hybrid equilibrium

We can again show numerically that when a mutant invades, it reaches a stable equilibrium value.

#### 6.3 Evolutionary route to asexuality

The partial derivatives of the fitness function in this model are:

$$\begin{cases} \dfrac{\partial w}{\partial \alpha'} = (1-\hat{\Phi})\dfrac{1+\sigma}{2}\dfrac{3\beta-1}{2} & \text{(26a)} \\ \dfrac{\partial w}{\partial \beta'} = (1-\hat{\Phi})\dfrac{1+\sigma}{2}\dfrac{3\alpha-1}{2} & \text{(26b)} \\ \dfrac{\partial w}{\partial \sigma'} = \dfrac{1-\hat{\Phi}}{2}\left(\alpha\beta + \dfrac{(1-\alpha)(1-\beta)}{2}\right) & \text{(26c)} \end{cases}$$

Eq. (26c) is trivially always positive: selection always favour more female-biased sex-ratios, as in previous models. Eq. (26a) and (26b) are positive if and only if $\beta > 1/3$ and $\alpha > 1/3$ respectively. This is always the case when a first mutant has been able to spread.

**If a first partially asexual mutant has spread, evolution should favour the eventual evolution of full asexuality.**

### RELAXING THE ABSENCE OF INTER-SPECIFIC COMPETITION

We illustrate here the quantitative influence of the competition parameter $\chi$ on the evolution of asexuality. As an example, we will take the case of the first scenario, *sperm fails to decondense*.

Assuming $\chi \neq 0$, then the dynamics of the hybrid population can be described with the equation:

$$\dot{n}_h = \theta + \left(\frac{1-\Phi}{2} - 2(n_h + 2\chi n_p)\right) n_h \tag{27}$$

This leads to a different equilibrium population size of the hybrid population.

When there is no inter-specific competition ($\chi = 0$), $\dot{n}_h \geq 0$ when $n_h$ tends to $0$. However, this is not the case anymore when $\chi > 0$. In this latter case, $\dot{n}_h$ can be $< 0$ when $n_h$ tends to $0$ if $\frac{1-\Phi}{2} - 4\chi n_p < -\frac{\theta}{n_h}$. Note that this can be true if and only if $\theta$ also tends to $0$ (when there a production of wild-type hybrids at each generation because of direct hybridization, this population cannot go extinct).

**When hybrids are rare, they may become extinct if the burden of inter-specific competition with parental species is greater than births obtained via (i) direct hybridization of parentals, and (ii) back-crossing of hybrids with parentals. Interestingly, because direct hybridization constantly produces hybrids from non-hybrid individuals, it tends to rescue hybrids from extinction; extinction can only really happen when there is no direct hybridization.**

In cases where the hybrid population survives and coexists with the parental populations, we can write the mutant hybrid population growth-rate as:

$$w = \frac{1+\sigma}{2}\left[\alpha\left(\hat{\Phi} + \left(1-\hat{\Phi}\right)\beta\right) + (1-\alpha)\frac{1}{2}\left(1-\hat{\Phi}\right)(1-\beta)\right] - 2(\hat{n}_h + 2\chi\hat{n}_p) \quad (28)$$

A first, partially clonal mutant can invade if and only if a mutant with phenotype $\{\alpha = 1, \beta = 0, \sigma = 0\}$ can (as this is the "best" clonal mutant possible). As such, we deduce that some partially clonal mutant can invade if the minimal invasion condition is met:

$$\frac{\hat{\Phi}}{2} > 2(\hat{n}_h + 2\chi\hat{n}_p) \quad (29)$$

This time we have $\dot{n}_h = 0$ that implies that $\frac{1-\hat{\Phi}}{2} + \frac{\theta}{\hat{n}_h} = 2(\hat{n}_h + 2\chi\hat{n}_p)$ instead of simply $2\hat{n}_h$. The minimal invasion condition when assuming $\chi \neq 0$ can thus be written as:

$$\frac{\hat{\Phi}}{2} > \frac{1-\hat{\Phi}}{2} + \frac{\theta}{\hat{n}_h} \quad (30)$$

Here, $\chi$ does not appear. This same invasion condition can indeed be written for $\chi = 0$. Inter-specific competition does not qualitatively change the outcome that at least some partially clonal mutant can invade if hybrid $\times$ hybrid matings are frequent enough, and direct hybridization rare enough.

To determine how $\chi$ quantitatively alters the results, we plot on Fig. (3) $\hat{\Phi} - \frac{1}{2} - \frac{\theta}{\hat{n}_h} = 0$ calculated numerically for different values of $\theta$, $c_h$ and $\hat{n}_p$. We determine graphically the areas where the mutant hybrids invade (when the parameters are such that $\hat{\Phi} - \frac{1}{2} - \frac{\theta}{\hat{n}_h} > 0$, which logically correspond for example to values of $\theta$ lower than the one such that the expression equals $0$). We see that invasion of mutants tend to happen at higher values of $c_h$ and lower values of $\theta$. This corresponds to the findings of the general analysis: invasion of mutants is promoted when hybrid females mate with hybrid males, and when hybridization between the parentals is rare. The intuitive explanation is that mutants not only compete with wild-type hybrids now, but also with parental populations. This leads to an increase of the overall number of competitors, and thus an increase of the overall death rate of mutant hybrids, making it more difficult for them to invade.

**Inter-specific competition tends to reduce the parameter space of mutant invasion as it increases the death rate of arising mutants. This effect increases with the size of the parental populations; when parental populations are small, they exert little competition on hybrids and thus the invasion of mutants is not strongly altered. Even though inter-specific competition tends to make it harder for mutants to invade, results are qualitatively similar for the whole range of $\chi$.**

The partial derivatives of $w$ are the same independently of whether or not we consider inter-specific competition. The routes towards asexuality are thus the same than before. Most importantly, this means that asexuality still fully evolves whenever the first mutation invades.

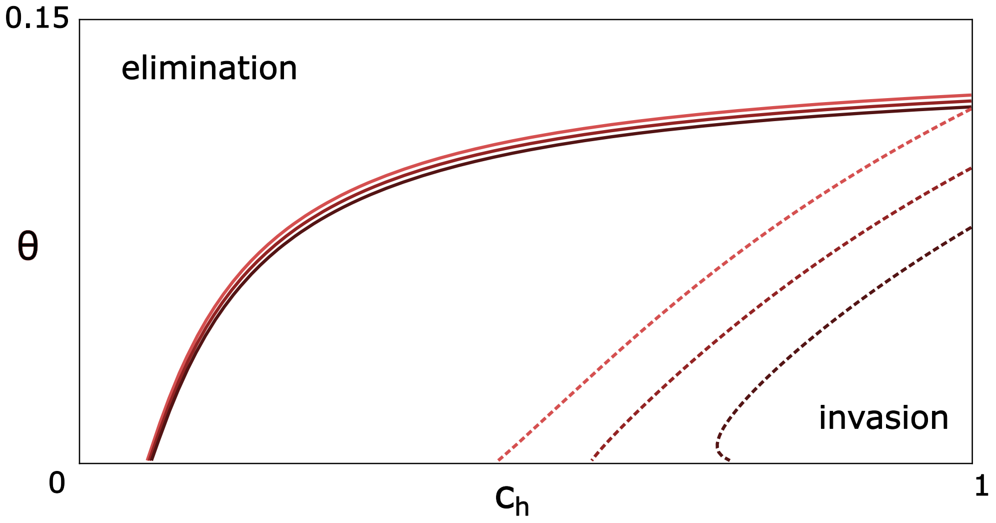


**Figure 3:** Mutant invasion $\{\theta, c_h\}$-parameter space for different values of $\chi$ and $\hat{n}_p$. Increasingly dark tones of red show $\chi = 0.1$, $\chi = 0.5$ and $\chi = 0.9$. Plain lines were calculated with $\hat{n}_p = 0.01$, while dotted lines were calculated with $\hat{n}_p = 0.1$. Mutation invasion happens for values of $\theta$ and $c_h$ below and to the right-hand side of the lines drawn.

### RELAXING THE COMPLETE SEXUALITY OF WILD-TYPE HYBRIDS

Throughout we assumed that wild-type hybrids are perfectly sexual: $\alpha_{wt} = \beta_{wt} = \sigma_{wt} = 0$.

However, empirical findings suggest that some F1 hybrids, though not readily asexual, do exhibit some partial asexuality. Particularly, there exists evidence of partial production of clonal eggs. Here, we illustrate how that would influence the outcome of the model, taking again the example of the first scenario, *sperm fails to decondense*. We assume in the following: $\beta_{wt} = \sigma_{wt} = 0$ and $\alpha_{wt} > 0$.

Under this new assumption, the dynamics of the wild-type hybrid population becomes:

$$\dot{n}_h = \theta + \left( \frac{(1 - \alpha_{wt})(1 - \Phi) + \alpha_{wt}\Phi}{2} - 2n_h \right) n_h \tag{31}$$

We can solve this differential equation and obtain numerically the stable equilibrium value of $\hat{n}_h$ and of $\hat{\Phi}$. Qualitatively, results are similar as when we assumed $\alpha_{wt} = 0$. We plot on Fig.(4) this new $\hat{n}_h$, as well as $\Delta\hat{n}_h$, the difference between $\hat{n}_h$ when $\alpha_{wt} > 0$ and when $\alpha_{wt} = 0$.

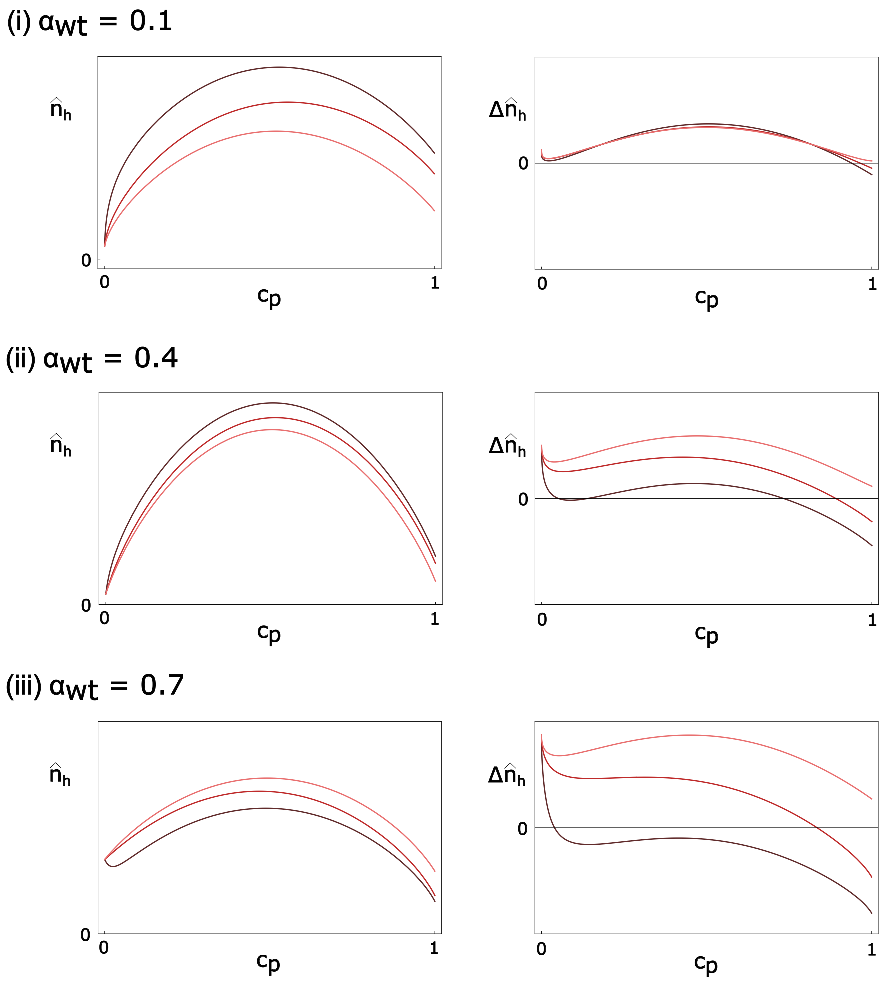


**Figure 4:** $\hat{n}_h$ (left-hand side column) and $\Delta\hat{n}_h$ (right-hand side column) for different values of $c_p$ (x-axis), $\alpha_{wt}$ and $c_h$. Darker tones of red indicate smaller values of $c_h$: $c_h = 0.9$, $c_h = 0.5$ and $c_h = 0.1$ from lighter to darker.

Generally, we see that partial asexuality tends to increase the wild-type hybrid population size ($\Delta\hat{n}_h > 0$). This may not always be the case however, especially when (i) $\alpha_{wt}$ is large, (ii) $c_h$ is small, and (iii) $c_p$ is large. The influence of $c_h$ is due to the fact that hybrids gain reproductive opportunities with hybrid males and lose reproductive opportunities with parental males. Thus, when hybrids tend to reproduce a lot with parental males (that is, when $c_h$ is small), partial asexuality is more of a burden for hybrid females and their population size actually decreases.

Invasion condition of mutants do not change by assuming that $\alpha_{wt} \neq 0$, as the phenotype of mutants is the same as before. However, numerically, changes in wild-type hybrid equilibrium population size should bring changes to the death rate of mutants (as they compete with wild-type hybrids for resources). That is, the parameter space that favours the invasion of mutants should numerically change with $\alpha_{wt}$. We show on Fig. (5) a comparison of the invasion conditions of mutants for $\alpha_{wt} = 0$, $\alpha_{wt} = 0.1$ and $\alpha_{wt} = 0.4$.

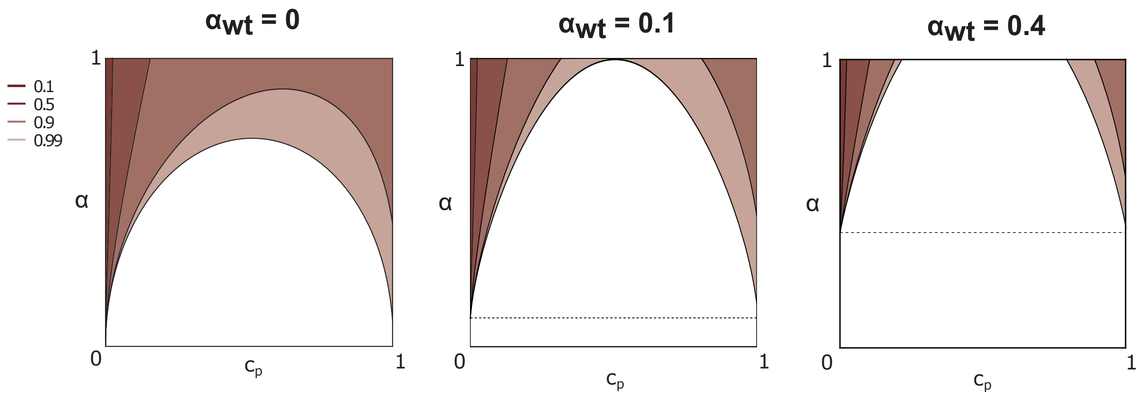


**Figure 5:** Minimal value of mutant rates of clonal egg production $\alpha$ necessary for a mutation to invade, for different value of $c_p$ (x-axis), $c_h$ and $\alpha_{wt}$ the wild-type rate of clonal egg production. Shaded areas correspond to areas where invasion can happen. Darker tones of red indicate lower values of $c_h$.

**We can see that as $\alpha_{wt}$ increases, it becomes more and more difficult for a mutant to invade. The spread of more-asexual mutations increasingly rely on extreme values of $c_p$ and high values of $c_h$. Note however that invasion conditions remain qualitatively similar in the three cases. Mutations have more trouble spreading because equilibrium population size of wild-type hybrids tend to increase. This increases the death rate of mutant hybrids by competition for resources with wild-type hybrids, hereby reducing their growth rate and invasion capacities.**

Importantly, we see that $\alpha < \alpha_{wt}$ can never invade: the route towards restoring sexuality in hybrids does not seem to be favourable. This is because asexuality is still favoured here; the reduction of its invasion space is only due to a reduction of the force selecting for it, not to a reversal of it. Also, we see that there is no point where the invasion of mutation is easier than when $\alpha_{wt} = 0$, despite equilibrium population size of wild-type hybrids being reduced by $\alpha_{wt} > 0$ in some cases. This is because these cases correspond to situations where asexuality is selected against (low $c_h$): in this parameter space, mutants do not invade.

The partial derivatives of $w$ are the same independently of whether or not we consider $\alpha_{wt} > 0$. The routes towards asexuality are thus the same than before. Most importantly, this means that asexuality still fully evolves whenever the first mutation invades.
